# Chromatin state barriers enforce an irreversible mammalian cell fate decision

**DOI:** 10.1101/2021.05.12.443709

**Authors:** M. Andrés Blanco, David B. Sykes, Lei Gu, Mengjun Wu, Ricardo Petroni, Rahul Karnik, Mathias Wawer, Joshua Rico, Haitao Li, William D. Jacobus, Ashwini Jambhekar, Sihem Cheloufi, Alexander Meissner, Konrad Hochedlinger, David T. Scadden, Yang Shi

## Abstract

Stem and progenitor cells have the capacity to balance self-renewal and differentiation. Hematopoietic myeloid progenitors replenish more than 25 billion terminally differentiated neutrophils every day under homeostatic conditions and can increase this output in response to stress or infection. At what point along the spectrum of maturation do progenitors lose capacity for self-renewal and become irreversibly committed to differentiation? Using a system of conditional myeloid development that can be toggled between self-renewal and differentiation, we interrogated determinants of this ‘point of no return’ in differentiation commitment. Irreversible commitment is due primarily to loss of open regulatory site access and disruption of a positive feedback transcription factor activation loop. Restoration of the transcription factor feedback loop extends the window of cell plasticity and alters the point of no return. These findings demonstrate how the chromatin state enforces and perpetuates cell fate and identifies potential avenues for manipulating cell identity.

**Highlights:**

- There exists a point of irreversible commitment in granulocytic differentiation
- Chromatin state dynamics establish the transition from self-renewal to differentiation commitment
- Reduced chromatin accessibility underlies an irreversible loss of regulatory site access
- Restoration of a transcription factor feedback loop alters the differentiation commitment point

## Introduction

Differentiation is generally understood to be a process whereby a stem or progenitor cell divides, matures, and eventually exits the cell cycle as a ‘terminally’ differentiated effector cell. An intriguing and fundamental aspect of differentiation is its unidirectionality; with rare exceptions^1–4^, mammalian cells do not revert or de-differentiate under physiological conditions^5,6^. This suggests that cell circuits embed critical stages of cell fate commitment, beyond which the differentiation program is irreversible. While the concept of differentiation commitment is well-appreciated, the mechanistic basis of this process is not understood. What are the molecular “locking” factors that enforce cell fate along the pathway from progenitor to effector cell? This question has significant biomedical ramifications, as successful manipulation of cellular identity is critical for tissue regeneration and pro-differentiation-based cancer therapies.

Several hypotheses seek to explain the irreversibility of differentiation. Epigenomic analyses of the induced differentiation of pluripotent cells offer correlative support to the importance of chromatin dynamics and describe a program in which lineage-specific transcription factors (TFs) are induced, and then reinforced by stable, chromatin-based silencing of pluripotency gene expression programs^7–9^. Proposed silencing mechanisms^10–12^ include DNA methylation and repressive histone modifications such as H3K27me3 and H3K9me3. These modifications are antagonistic to transcription and are self-perpetuating^13–16^, offering a means of permanently silencing transcription in the absence of continued stimulus. Studies in reprogramming also indirectly support this hypothesis. By exogenously expressing four key TFs, mature cells can be reverted to pluripotent cells^17^. Importantly, three of these four factors – OCT3/4, SOX2, and KLF4 – are “pioneer” TFs, denoting that they can bind their DNA motifs even in the presence of nucleosomes and may override repressive chromatin states^18,19^. However, most TFs are not pioneer factors, allowing repressive chromatin states to enforce cellular identity under normal conditions.

While compelling, these hypotheses have been difficult to test. Time-course profiling of differentiation programs captures features that correlate with maturation, though it is not clear which of these features are causal in preventing reversion of the program^20^. Furthermore, the large-scale profiling studies that generate such models most commonly focus on the differentiation of embryonic stem cells^7–9,11,21^. These models may be applicable for cell fate decisions in early embryogenesis, but it is not known whether they also apply to adult stem and progenitor cell developmental programs.

To understand mammalian cell differentiation commitment, we have employed a system of conditional myeloid progenitor cell differentiation arrest that can be released from differentiation blockade under tight temporal control^22^. Using this system, we identified a critical differentiation commitment window, beyond which the cells are incapable of returning to the progenitor state. By performing epigenomic profiling before, during, and after this window of commitment, we identified the chromatin state and transcriptomic processes that determine differentiation commitment.

Unexpectedly, we did not observe promoter DNA methylation or H3K27me3 as contributors of differentiation commitment. Instead, the most striking dynamics in this program involved chromatin remodeling, enhancer activation, and transcription factor usage. Differentiation was accompanied by a progressive global loss of accessible chromatin and concomitant stable silencing of progenitor state TFs. Importantly, ectopic expression of the transcription factors *Meis1* and *Pbx1* significantly extends the duration of cells along the differentiation pathway before they are irreversibly committed to terminal differentiation. Here we propose a parsimonious model in which the loss of access to active regulatory sites is sufficient to disable TF-driven re-acquisition of the progenitor state. This model highlights the essential interaction between TFs and chromatin and ascribes a new role to enhancers in cellular identity – the ability to act as barriers to de-differentiation.

## Results

### Myeloid differentiation is initiated by repression of progenitor state transcriptional programs

Studies of myeloid cell differentiation commonly rely on sorted populations of primary cells, which are limited in abundance, or on cancer cell lines, which often differentiate aberrantly and in response to non-physiologic stimuli. To study the mechanism of normal differentiation commitment, we used a system that provides an abundant supply of diploid progenitor cells capable of synchronous and terminal differentiation. The ER-HOXA9 and ER-HOXB8 models of conditional differentiation arrest offer these benefits^22,23^. In the presence of estrogen (beta-estradiol, E2), the ER-HOXA9 transcription factor is active and prevents differentiation beyond the granulocyte-monocyte precursor (GMP) stage, allowing for unlimited expansion. Withdrawal of estrogen rapidly inactivates the ER-HOXA9 protein via nuclear export and cytoplasmic sequestration. Following ER-HOXA9 inactivation, cells synchronously progress through normal myeloid differentiation over ∼5 days to fully mature neutrophils and (less commonly) monocytes.

In utilizing HOXA9 (or HOXB8) to arrest differentiation, this model takes advantage of our understanding of normal hematopoietic development. Progenitors must downregulate *Hoxa9* expression as they commit to becoming mature monocytes and neutrophils. HOXA9 is a well-appreciated regulator of myeloid cell fate and mice lacking *Hoxa9* have reductions in circulating white blood cell counts, myeloid colony forming ability, and responsiveness to G-CSF^24^. The dysregulated persistence of *Hoxa9* expression promotes GMP differentiation arrest and is a key oncogenic event seen in >70% of human AML^25^.

These particular ER-HOXA9 GMPs were derived from adult bone marrow of a Lysozyme-GFP knock-in mouse^26^. In this context, expression of GFP acts as an endogenous reporter of differentiation status, as only maturing neutrophils activate expression of the secondary granule protein lysozyme.

To investigate the dynamics of differentiation, we analyzed gene expression (RNA-seq) at 10 time points across the 120-hour program. The differentiation program proceeded in graded fashion, with progressive and continuous global trends in gene expression (Figures 1A and Figure S1A). We identified six main patterns of gene expression that differed primarily in the kinetics of induction or repression (Figure 1B).

**Figure 1.**
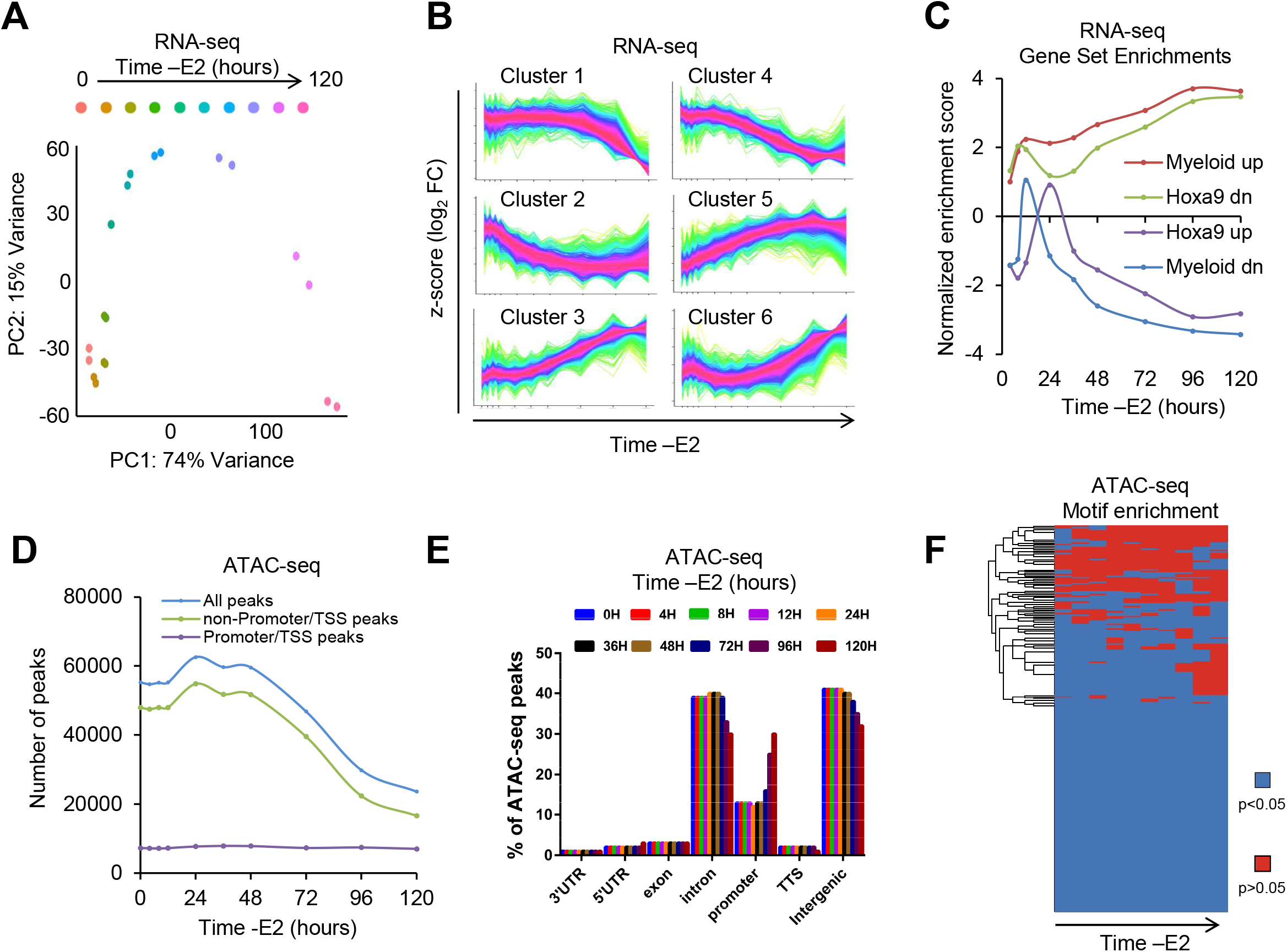
Transcriptomic and ATAC-seq profiling of the myeloid differentiation program. A) Principal component analysis (PCA) of RNA-sequencing at 10 time points in biological duplicate following inactivation of ER-HOXA9 (upon removal of estradiol) over 120 hours. B) Identification of six main clusters of genes with distinct expression dynamics over the differentiation program. Individual transcripts in each cluster are colored according to degree of correlation to the cluster trend. C) Plot of GSEA normalized enrichment scores (NES) of myeloid differentiation gene sets over the differentiation time-course. D) Plot of number of ATAC-seq peaks in ER-HOXA9 cells over the differentiation program. E) Distribution of ATAC-seq peaks at each time point according to genome annotation. F) Clustering of DNA motifs by degree of enrichment in ATAC-seq peaks of open chromatin at each time point. RNA-seq and ATAC-seq data represent the average of experiments performed in biological duplicate at each time point.

Comparing the 0-hour (progenitor state) time point to later time points, Gene Set Enrichment Analysis (GSEA)^27,28^ revealed that the interferon-γ response and TNF-α pathway were strongly induced throughout the time-course, and dramatic cell cycle repression and downregulation of Toll-like receptor X signaling emerged near the end of the program (Figures S1B and S1C). At the final time point the most highly enriched gene sets (of 3,477) were (1) Myeloid Development and (2) Genes Downregulated Upon Overexpression of Hoxa9 and Meis1 (Figure 1C and S1B, and Supplementary Table 1), confirming the full execution of the differentiation program. Interestingly, enrichment of these signatures over time showed unstable kinetics over the first 24-36 hours, after which they progressed consistently (Figure S1C (left panel)).

Analyses of TF and miRNA binding motif enrichment in promoters of differentially expressed genes identified the loss of MYC activity as an early event, induction of PU.1 and ELF1 targets as intermediate events, and loss of E2F1 as a terminal event (Figure S1C, right panel). On a global level, concerted downregulation of active TF/miRNA programs occurs prior to the strongest upregulation of a new set of TF/miRNA programs, suggesting that progenitor gene expression programs are shut down prior to induction of differentiation programs (Figure S1D).

ATAC-seq was performed to investigate epigenomic dynamics accompanying ER-HOXA9 differentiation. While patterns of chromatin accessibility evolved as cells differentiated (Figures S2A and S2C), the total amount of open chromatin appeared relatively constant in the first two days of the differentiation program. However, the number of ATAC-seq peaks dropped precipitously from 48 hours until terminal differentiation, with fully mature cells harboring less than half the amount of open chromatin as progenitor state cells (Figure 1D). Relatively few new regions of chromatin opened as cells differentiated, with the majority (75%) of accessible chromatin in differentiated cells already being open at the progenitor state (Figure S2B).

Concomitant with accessibility losses, the genomic distribution of ATAC-seq peaks showed a moderate relative shift from introns and intergenic regions to promote/TSS regions (Figure 1E). Interestingly, this shift did not reflect an increase in global promoter accessibility, as the absolute number of promoter/TSS peaks was constant through the time course, but instead resulted from the large drop in the absolute number of intron/intergenic peaks starting around 48 hours. (Figures 1D and 1E). We next considered motif enrichment in areas of accessible chromatin. Progenitor state open chromatin was enriched for binding sites of numerous TFs expressed both in the progenitor and the mature state (Figures 1F and S2D and Supplementary Table 2). Motifs enriched only at the early timepoints included binding sites for notable progenitor state TFs such as c-MYC, several GATA factors, and multiple HOX TFs. The small number of motifs enriched only in mature cells included binding sites for the neutrophil master regulators GFI1b, HIF1A, and NF-κB, suggesting the induction of these transcriptional programs late in the differentiation program.

Collectively, these data suggest that in the progenitor state, the chromatin landscape of ER-HOXA9 cells is already largely accessible to drivers of the differentiation program, but that progenitor state transcriptional programs must be silenced prior to the execution of terminal differentiation and concomitant global losses in chromatin accessibility.

### Identification of an irreversible “point of no return” in the myeloid differentiation program

In normal mammalian cells, terminal differentiation is thought to be irreversible. However, our transcriptional and epigenomic profiling did not identify any obvious inflection points or rapidly induced or lost gene expression programs that would suggest a critical commitment point in the program. To test whether the myeloid differentiation program is irreversible throughout, we temporarily inactivated ER-HOXA9 by estrogen withdrawal (“-E2 pretreatment”) for varying periods of time before reactivating it by the reintroduction of estrogen followed by monitoring the resulting extent of differentiation (Figure 2A).

**Figure 2.**
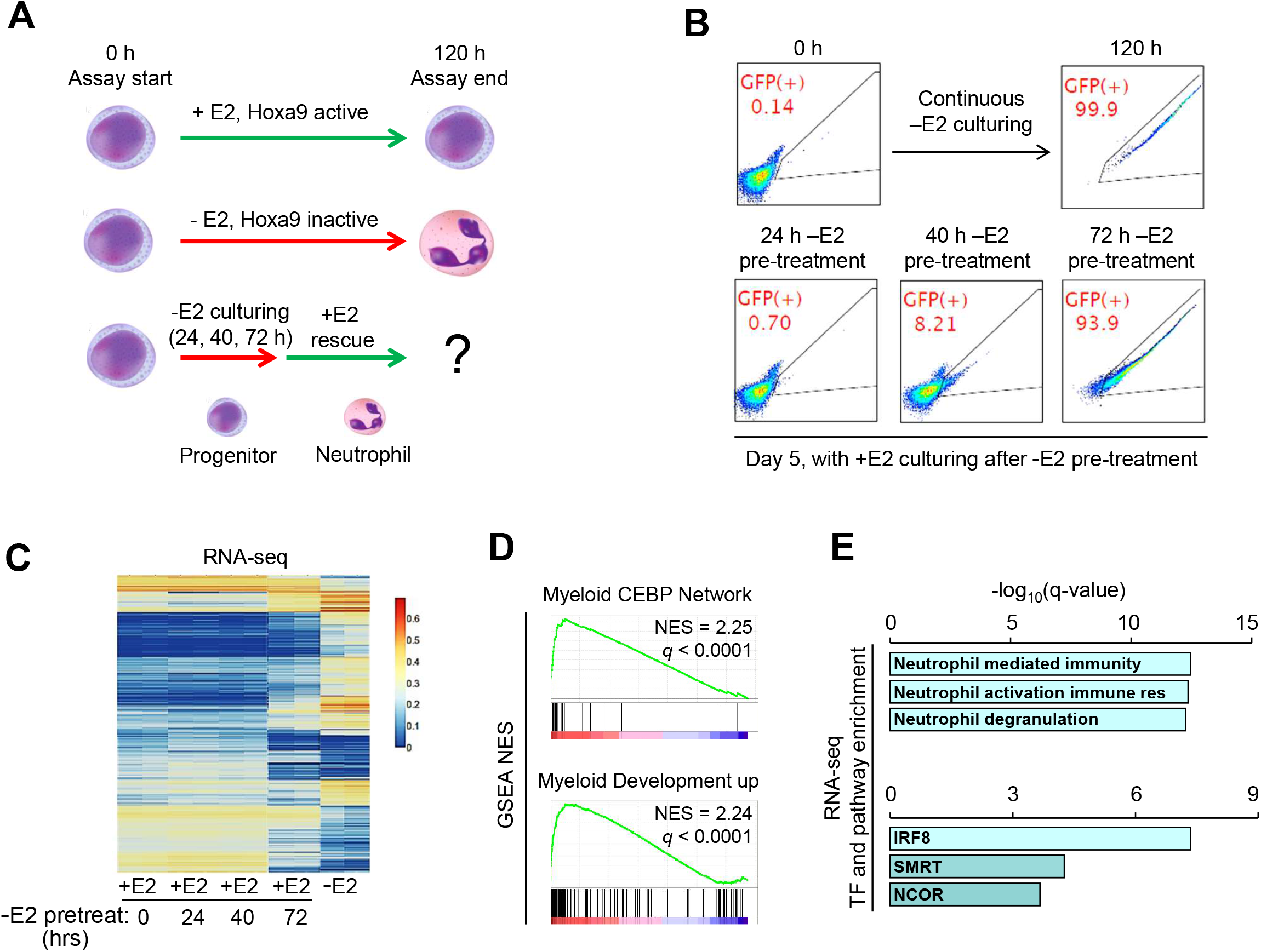
Identification of differentiation commitment “point of no return”. A) Diagram of scheme in which ER-HOXA9 is temporarily inactivated by estrogen withdrawal (-E2) for a period of time (“-E2 pretreatment”) before reactivation via E2 add-back and cell harvest for phenotypic and epigenomic analyses. B) Flow cytometry of GFP differentiation reporter in ER-HOXA9 cells in the progenitor state (0 h –E2), terminally differentiated state (120 h –E2), or in cells receiving -E2 pretreatments followed by E2 add-back to reactivate ER-HOXA9. Cells cultured with 24-hour or 40-hour -E2 pretreatment return to baseline after E2 add-back, whereas cells cultured with 72 -E2 pretreatment cannot be reverted to baseline by subsequent Hoxa9 reactivation. Images shown are representatives from biological triplicates. C) RNA-seq of cells cultured +E2 (progenitor state), 120 hours -E2 (terminally differentiated state) or +E2 after -E2 pretreatments. D) GSEA indicates that cells receiving 72-hour -E2 pretreatment stably maintain enrichment of the myeloid development maturation gene expression programs after ER-HOXA9 reactivation. E) GO biological process enrichment (top) and ChIP-seq-based transcription factor target enrichment (bottom) in the 200 genes most overexpressed in cells receiving 72-hour vs. 0-hour -E2 pretreatment.

According to our initial RNA-seq profiling (Figures 1A-1C), inactivation of ER-HOXA9 for 24 or 40 hours initiated early to intermediate steps of the differentiation program. However, we found that these early events in differentiation were reversible; reactivation of ER-HOXA9 overrode these changes and reverted cells back to (or very close to) the starting progenitor state (Figure 2B). In contrast, after inactivating ER-HOXA9 for 72 hours or more, the cells were stably committed to the differentiation program and could not be returned to the progenitor state upon ER-HOXA9 reactivation (Figure 2B); the cells had achieved a “point of no return” and inexorably progressed to terminal differentiation.

To confirm that this differentiation point of no return was not specific to a single ER-HOXA9 clone, we repeated the estrogen withdrawal experiments in other ER-HOXA9 clones as well as in the parallel iteration of the system immortalized by ER-HOXB8 expression^23^. While the exact time of irreversibility varied slightly, all versions of this system demonstrated a clear commitment point from which only rare cells could escape – even if the assay was extended to seven days - perhaps due to their lagging behind the rest of the differentiating population (Figures S3A and S3B).

To complement these reporter-based differentiation assays, we expanded our analysis to the transcriptomic level. In pre-commitment cells (24 or 40 hours -E2), reactivation of ER-HOXA9 successfully re-established a pattern of gene expression that was nearly indistinguishable from the transcriptome of naïve +E2 progenitor cells (Figure 2C). In contrast, in cells withdrawn from E2 for 72 hours, reactivation of ER-HOXA9 was unable to reverse the global induction of differentiation. The transcriptomes of these cells remained markedly different from progenitors and were strongly enriched for myeloid development gene signatures and activated neutrophil gene ontologies (Figures 2C-2E). By 72 hours -E2, the cells had passed a key commitment point, beyond which their phenotype was maintained in the absence of the originating stimulus. While not all cells in these samples proceeded to the very final stages of terminal differentiation (as seen in the 120-hour -E2 samples), we reasoned that the transcriptional changes maintained after 72-hours out of estrogen may represent the processes most critical for preventing reversion of the cells back to the progenitor state.

### Promoter DNA methylation and H3K27me3 silencing do not influence myeloid differentiation commitment

How was the differentiation gene expression program maintained following reactivation of ER-HOXA9? We first investigated the roles of DNA 5mC methylation and PRC2-mediated H3K27me3, which are repressive chromatin modifications that can be self-propagated, and during ESC differentiation often localize to genomic loci that are stably silenced in the chosen lineage^7–9^. Accordingly, we hypothesized that DNA methylation and/or H3K27me3 may facilitate commitment to myeloid differentiation.

We performed Reduced Representation Bisulfite Sequencing (RRBS)^29^ of 5mC as well as ChIP-seq of H3K27me3 on cells in the progenitor state (+E2), the terminally differentiated state (-E2 for 120 hours) and after withdrawing E2 for varying time periods before ER-HOXA9 reactivation. Unexpectedly, very few promoters showed differentially methylated regions (DMRs). In fact, the methylomes were so similar that biological replicates did not always cluster together (Figure S3C), and the few promoter DMRs that were observed did not correlate with transcriptional silencing (Figure S3D). Similar results were found when considering DNA methylation in H3K27ac ChIP-seq peaks (discussed below) marking putative active regulatory sites beyond promoters (Figure S3E). This suggests that DNA CpG methylation at promoters and regulatory elements does not strongly influence the transcriptional dynamics in this differentiation program.

We next examined H3K27 trimethylation. In progenitor cells, the H3K27me3 peaks were enriched at genes regulating early embryonic developmental programs, with dramatic enrichment (p<1×10^−102^) for homeodomain-containing genes (Supplementary Table 3). However, global H3K27me3 enrichment patterns did not change appreciably in differentiated cells (Supplementary Table 3 and Figures S4A-S4C), and H3K27me3-marked genes were not enriched for GO categories relevant to myeloid differentiation in any of the samples (data not shown). Furthermore, loss or gain of H3K27me3 from the progenitor to the differentiated state did not correlate with gene expression changes (Figure S4D). More specifically, genes comprising the Myeloid Development expression signature, the majority of which are highly differentially expressed over the differentiation program, had no appreciable changes in H3K27me3 dynamics (Figure S4E). We therefore conclude that, similarly to DNA promoter CpG methylation, H3K27me3 dynamics are unlikely to drive myeloid progenitor differentiation commitment.

### Chromatin remodeling is highly dynamic during differentiation

Given that we did not identify a role for DNA promoter CpG methylation or H3K27me3 in enforcing differentiation commitment and considering the dramatic decline in chromatin accessibility observed via ATAC-seq (Figure 1D), we investigated whether chromatin remodeling and regulatory site access dynamics could help lock cells into the maturation program. We compared ATAC-seq signatures between progenitor cells, mature cells, and in cells following the same -E2 pretreatment/add-back scheme described above. We also performed ChIP-seq of H3K27ac and H3K4me3 on matching samples to help map regulatory elements. ATAC-seq trends were highly dynamic, with unbiased clustering identifying seven main patterns of loci accessibility over the maturation program (Figure S5A). Progenitor cells were again observed to have roughly twice as many peaks as the terminally differentiated cells (Figures 3A and 3B). Intriguingly, longer -E2 pretreatments led to greater irreversible losses of ATAC-seq peaks up to the 56-hour -E2 pretreatment sample, suggesting significant epigenetic “memory” of the low chromatin accessibility state even after ER-HOXA9 reactivation.

**Figure 3.**
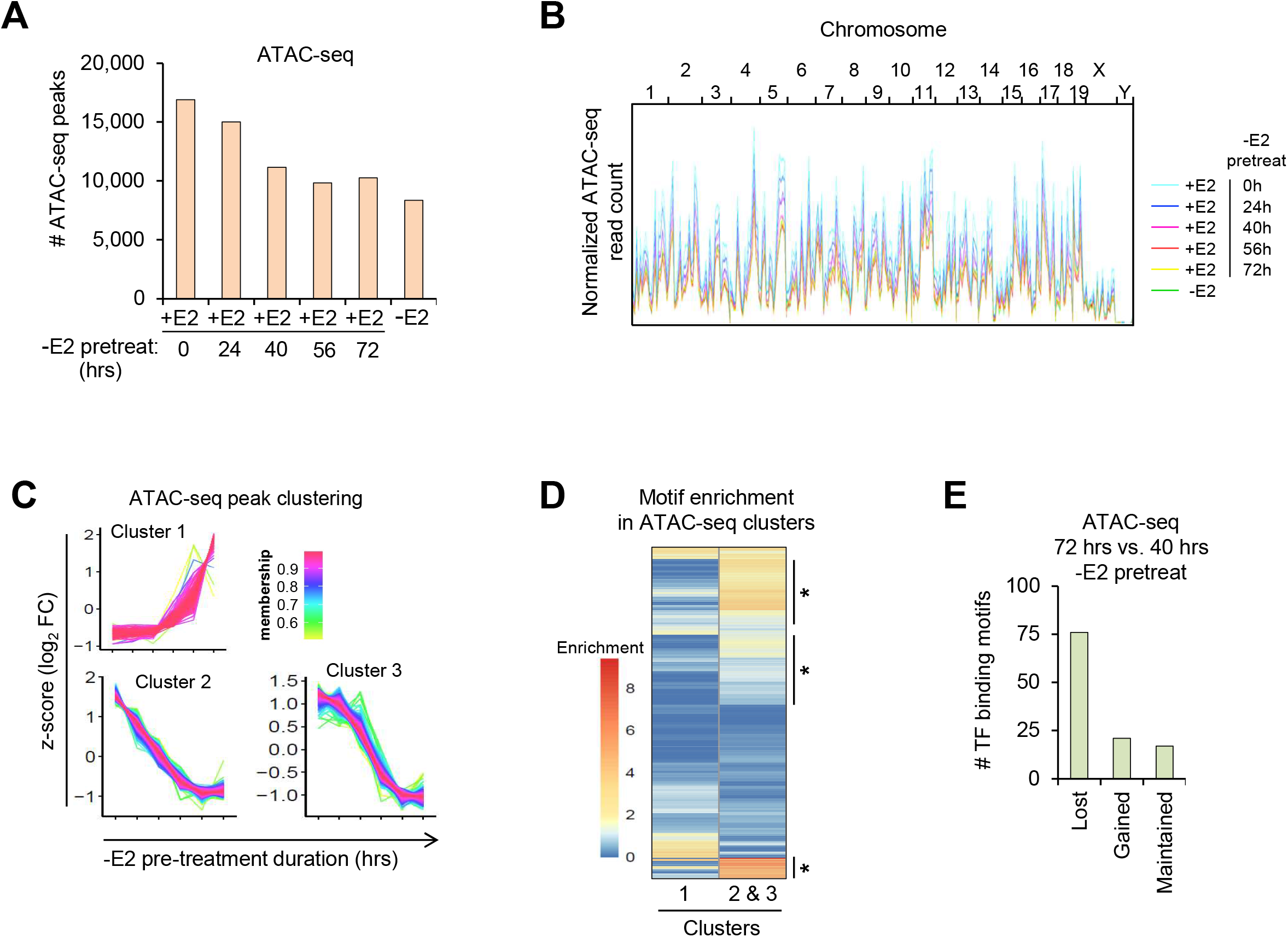
Chromatin remodeling dynamics in differentiation commitment. A) Number of ATAC-seq peaks in ER-HOXA9 cells cultured +E2 (progenitor state), -E2 for 120 hours (terminally differentiated state), or +E2 after -E2 pretreatments of varying duration. B) Normalized ATAC-seq read counts of samples across all chromosomes. C) Identified clusters of genomic loci undergoing the most dramatic ATAC-seq dynamics in samples receiving 40-hour compared to 72-hour -E2 pretreatment prior to E2 add-back (representing samples that did or did not pass the commitment point, respectively). D) TF binding motif enrichment in ATAC-seq peaks found in clusters shown in (C). Heat map is colored by motif enrichment score in each sample. Clusters of motifs that are enriched in 72-hour -E2 pretreatments are noted with lines and asterisks. E) Quantification of binding motif enrichment trends in heat map shown in (D). ATAC-seq was performed in biological duplicate at all time-points and results represent the average at each time-point.

We compared chromatin accessibility in the populations receiving -E2 pretreatments of 40 hours or 72 hours. Here, differentiation was initiated in both samples (Figures 1C and 2B), but only the 40-hour sample could be reverted to the progenitor state by ER-HOXA9 reactivation. Three main clusters of accessible chromatin dynamics were identified when comparing these two samples: two regions that lost accessibility, and one region that gained accessibility (Figure 3C). Motif enrichment of these genomic regions showed that many TF binding sites become irreversibly depleted from accessible chromatin in cells 72-hours out of estrogen as compared to 40-hours, while few motifs gained enrichment (Figures 3D and 3E).

TFs with binding motifs no longer enriched in open chromatin after the commitment point include many notable drivers of the progenitor/proliferation program, such as RUNX1 and 2, ERG, E2F1, and several ETS factors. Concordantly, loci of canonical transcriptional targets of these TFs, such as *Bcl2* and *Hmga2* (RUNX1^30^), *Cdkn1a* and *Rfc1* (E2F1^27,31,32^), and *Gata2* and *Arid3a* (ERG^33^), had regions of open chromatin in the +E2 and 40 h –E2 pretreatment samples, but not in the –E2 or 72 h –E2 pretreatment samples (Figure S5B). Thus, a stable, irreversible reduction of TF access to chromatin distinguishes cells that pass beyond the differentiation commitment point.

Altogether, our findings suggest a model in which transcriptional repression of pro-stemness TFs is followed by losses in chromatin accessibility at their target loci, blocking their access to chromatin upon re-expression. This model offers a parsimonious explanation for why long-term repressive epigenetic modifications such as H3K27me3 and promoter DNA methylation may not be a pre-requisite for differentiation commitment in adult progenitor cells.

### Identification of ER-HOXA9 direct target genes

To test our model, we focused on the most critical pro-stemness driver TF in this system, ER-HOXA9 itself. Our model predicts that, once the differentiation program is initiated, ER-HOXA9 can access its progenitor-state binding sites only if re-activated prior to the commitment point. To identify direct ER-HOXA9 binding sites, we turned to ChIP-seq and re-established new model cell lines with an ER-HOXA9 (or WT HOXA9) construct harboring both V5 (N-terminal) and AM (C-terminal) epitope tags. We evaluated the V5 and AM antibodies in ChIPs of the wild type (non-ER fusion) V5-HOX9-AM protein in progenitor cells. While V5 ChIPs yielded more peaks than AM ChIPs (17,214 vs. 6,876, respectively), other results were highly consistent. V5 and AM ChIPs produced extremely similar global enrichment patterns with expected mapping to intronic and distal regions^34,35^, and had highly overlapping gene targets, GO enrichments, and motif enrichments (Figures S6A-S6F and Supplementary Table 4). However, in ChIPs of the ER-HOXA9 fusion protein, the V5 antibody appeared to have outperformed the AM antibody and closely recapitulated binding patterns of the wild type protein, albeit with fewer peaks overall (Figures 4A-4D). Accordingly, we used the V5 tag for all subsequent ChIP experiments.

**Figure 4.**
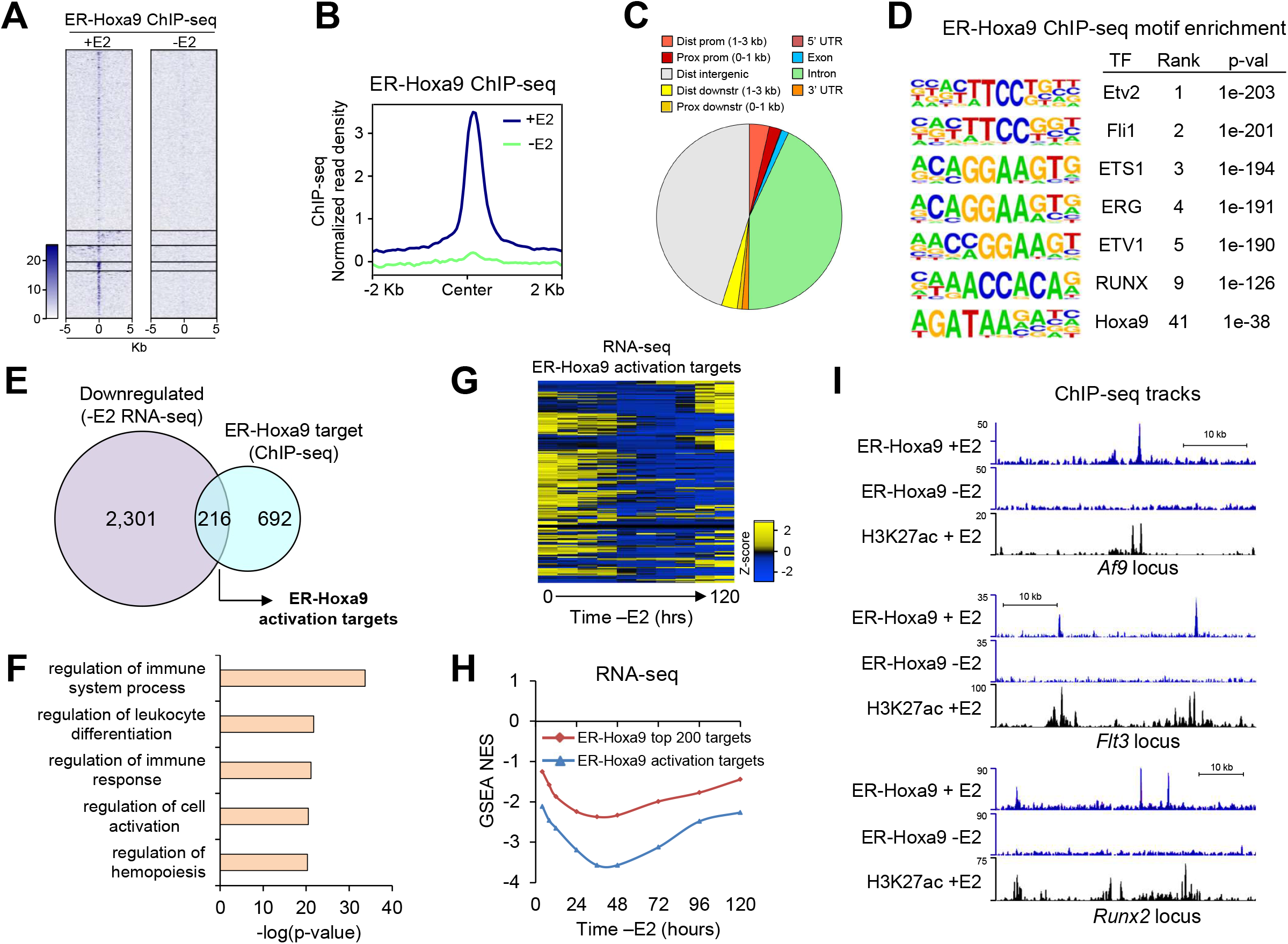
Identification of ER-HOXA9 direct targets. A) Heat map of merged peak regions from V5 ChIP-seq of V5-ER-HOXA9-AM cultured +E2 (nuclear V5-ER-HOXA9-AM) or 96 hours -E2 (cytoplasmic V5-ER-HOXA9-AM). Regions shown are +/- 5Kb from the peak center. B) Plot of average V5-ER-HOXA9-AM ChIP-seq normalized read density from +E2 and -E2 samples. Window shown is +/- 2 Kb from center of all +E2 peak regions. C) Distribution of +E2 V5-ER-HOXA9-AM peaks by genome annotation. D) MEME enrichment of known DNA motifs within peaks from +E2 V5-ER-HOXA9-AM ChIP-seq. E) Overlap of genes that are transcriptionally downregulated within 48 hours of -E2 treatment and genes with proximal +E2 V5-ER-HOXA9-AM ChIP-seq peaks. These are considered putative direct activation targets of ER-HOXA9. F) Top 5 GO biological processes enriched in ER-HOXA9 direct activation targets. G) Heat map of transcriptional dynamics of ER-HOXA9 activation targets over the 120-hour - E2 differentiation program. H) GSEA normalized enrichment scores (NES) of V5-ER-HOXA9-AM top 200 gene targets (blue) and 216 direct activation targets (red) tested for enrichment in RNA-seq of genes ranked by expression at each -E2 timepoint relative to 0-hour (+E2) timepoint. I) V5-ER-HOXA9-AM and H3K27ac ChIP-seq tracks at loci three direct activation targets of ER-HOXA9 from cells cultured +E2 or -E2.

We sought to identify the direct regulatory targets of ER-HOXA9. Progenitor state (+E2) V5-ER-HOXA9-AM ChIP-seq yielded 1,417 peaks that mapped to 908 genes (Supplementary Table 5), a subset of which we expect are functionally regulated by ER-HOXA9. Consistent with this, the set of the top 200 putative ER-HOXA9 target genes were rapidly globally downregulated as cells differentiated, reaching maximum repression by 36 hours -E2 (Figures 4G and 4H).

We defined the putative ER-HOXA9 direct, functional activation targets as genes bound by ER-HOXA9 in progenitor cells that are significantly downregulated at any point in the first 48 hours of differentiation. This yielded 216 genes that are highly enriched for GO categories involving the immune response and hematopoiesis (Figures 4E-4H and Supplementary Table 5). Intriguingly, among these targets are several critical transcriptional regulators known to drive the progenitor/proliferative state and contribute to acute myeloid leukemia (AML), including *Runx2, Meis1, Gfi1b*, and *Af9*, as well as other AML oncogenes such as *Flt3* and *Msi2* (Figure 4I). The direct regulation of other master TFs by ER-HOXA9 suggests that orchestration of progenitor state identity may utilize a hierarchical TF control model^36^, in which a very small number of master regulators have disproportionately large control of the gene regulatory network. Importantly, this model predicts that TFs at or near the top of the hierarchy have a very small number of most critical regulatory targets (i.e., a few other TFs).

We then investigated whether ER-HOXA9 can access its binding sites at progenitor state genes when reactivated in cells that have passed the differentiation commitment point. We induced differentiation by withdrawing E2 for 72 hours and then reactivated ER-HOXA9 and collected samples for ChIP-seq. Consistent with significant loss of chromatin access, reactivated ER-HOXA9 yielded markedly reduced ChIP-seq signal (Figures 5A-5B) and was only bound to 240 of its original 1,417 progenitor state binding sites (16.9%), as well as to 187 new binding sites (Figure 5C and Supplementary Table 5). Critically, only 32 ER-HOXA9 direct activation target genes were bound by reactivated ER-HOXA9 (Supplementary Table 5), and these did not include any of the aforementioned master TFs. Consequently, despite ER-HOXA9 reactivation, these essential drivers of the progenitor state remained irreversibly downregulated after the commitment point (Figure 5D). This is in line with the observation that the functional activation targets of ER-HOXA9 were among the earliest genes to be silenced upon E2 withdrawal (Figure 4G and 4H), and hence the least likely to retain open chromatin late in the differentiation program. Of note, several of the original target loci that reactivated ER-HOXA9 could bind to included genes that are highly upregulated in differentiated cells (*Lysozyme, Mmp8, Cleck4n, Ccl4*, and others).

**Figure 5.**
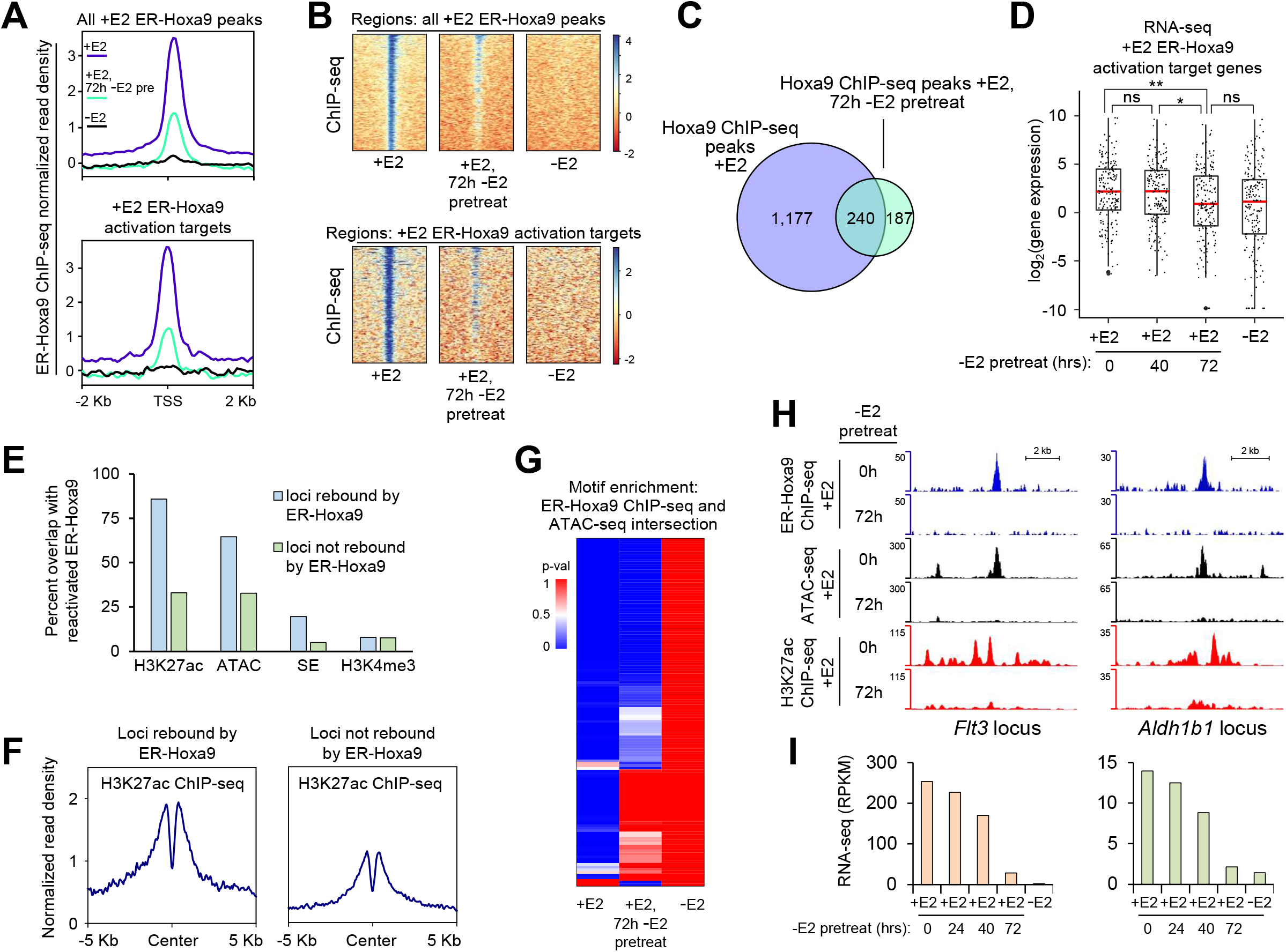
Integrative epigenomics analysis of ER-HOXA9 binding patterns in pre- and post-commitment cells. A) Plots of normalized read density from ChIP-seq of V5-ER-HOXA9-AM in cells cultured +E2 (purple), -E2 (black), and +E2 after 72-hour -E2 pretreatment (light green). Top plot is averaged over all +E2 V5-ER-HOXA9-AM peaks, bottom plot is averaged over the +E2 direct activation targets of V5-ER-HOXA9-AM. B) Heat map visualization of ChIP-seq data shown in (A). C) Overlap of V5-ER-HOXA9-AM peaks from cells cultured +E2 or +E2 after 72-hour -E2 pretreatment. D) Box and whisker plot representation of expression of V5-ER-HOXA9-AM activation target genes in cells cultured +E2, 120 hours -E2, or +E2 after -E2 pretreatments. Red line indicates median, and whiskers represent +/- 1.5*IQR (IQR = interquartile range). **\***t-test p < 0.05; **t-test p < 0.01. ns = not significant. E) Percent of V5-ER-HOXA9-AM peaks overlapping an H3K27ac peak, ATAC-seq peak, superenhancer, or H3K4me3 peak. V5-ER-HOXA9-AM peaks are split into groups of +E2 binding sites that ER-HOXA9 can (light blue) or cannot (light green) re-bind when reactivated after 72-hour -E2 pretreatment. F) Plots of H3K27ac ChIP-seq average read density in loci that ER-HOXA9 can (left) or cannot (right) re-bind when reactivated after 72-hour -E2 pretreatment. G) Heat map of DNA motif enrichments in the intersection of ATAC-seq and V5-ER-HOXA9-AM ChIP-seq peaks in cells cultured +E2, 120 hours -E2, or +E2 after 72-hour -E2 pretreatment. Heat map is colored according to p-value of motif enrichment. H) Tracks of V5-ER-HOXA9-AM and H3K27ac ChIP-seq along and matching ATAC-seq at the *Flt3* (left) and *Aldh1b1* (right) loci in cells cultured +E2, with or without 72-hour -E2 pretreatment. I) Expression of *Flt3* (left) and *Aldh1b1* (right) in cells cultured +E2, 120 hours -E2, or +E2 after -E2 pretreatments.

### Chromatin state barriers restrict ER-HOXA9 activity in post-commitment cells

We next determined the features of chromatin that precluded ER-HOXA9 binding after the commitment point. We separated progenitor state bindings sites into those that ER-HOXA9 could (n=240) or could not (n=1,117) re-bind to when reactivated after the commitment point. We then compared these sites for areas of overlap with our matching ATAC-seq, H3K27ac, H3K4me3, and H3K27me3 ChIP-seq datasets from post-commitment cells. In support of our model, the loci at which ER-HOXA9 failed to re-bind had dramatic aggregate reductions in ATAC-seq peaks, H3K27ac peaks, and super-enhancers compared to the loci with successfully re-bound ER-HOXA9 (Figures 5E and 5F). Peaks of H3K27ac, in particular, overlapped with 86% of the re-bound ER-HOXA9 sites, but with just 33% of the loci where ER-HOXA9 failed to re-bind. In contrast, there were no significant differences between re-bound and non-re-bound loci in overlap with H3K4me3 or H3K27me3 (Figure 5E and data not shown). These results suggest that the progenitor state enhancer landscape must be maintained for ER-HOXA9 to successfully drive progenitor cell identity.

Finally, we investigated the sequence content of the most high-confidence ER-HOXA9 target sites by intersecting ER-HOXA9 ChIP-seq and ATAC-seq peaks and performing motif enrichment analyses. As expected, open chromatin (ATAC-seq peaks) intersecting ER-HOXA9 peaks in the progenitor state was highly enriched for binding sites of TFs involved in progenitor biology, including ETS factors, HOXA9 itself, RUNX, MEIS, PU.1, and others (Figure 5G and Supplementary Table 6). Strikingly, fewer than half of these motifs were enriched in open chromatin intersecting ER-HOXA9 peaks in post-commitment cells, with binding sites for key TFs such as MEIS1 and PU.1:IRF notably absent. Accordingly, the loci of several potentially critical ER-HOXA9 targets, such as *Flt3, Aldh1b1, Pkca* and *Angpt1*, were largely inaccessible in post-commitment cells (Figures 5H, 5I and data not shown). This suggests that, following differentiation commitment, most of the original ER-HOXA9 binding sites are closed and that those that remain open harbor fewer options for coordinated binding between ER-HOXA9 and other transcriptional drivers of the progenitor state. Ultimately, these events compromise the ability of ER-HOXA9 to re-activate its activation targets, many of which remained irreversibly epigenetically silenced (Figures 5H and 5I).

In summary, in the progenitor state, ER-HOXA9 binds open chromatin at numerous genes and functionally regulates a subset of these. Upon inactivation of ER-HOXA9, its direct targets are rapidly downregulated with initiation of the differentiation transcriptional program, and as chromatin accessibility globally and progressively decreases, access to regulatory sites of these ER-HOXA9 targets declines. By 72 hours, most of the original binding sites are closed, and if ER-HOXA9 is re-activated, it can bind to only a small minority of sites that are still accessible. These do not include its most critical progenitor state regulatory targets; instead, these sites correspond to the small subset of genes that were initially bound by ER-HOXA9, but seem to play later roles in driving differentiation, and thus have maintained open chromatin status (Figure 7).

### Enforced *Meis1* expression alters the point of the no return by increasing cellular plasticity

While ER-HOXA9 binds at ∼1,000 genes (Supplementary Table 5), it appears to directly regulate approximately one quarter of these. In this subset are a handful of other transcriptional regulators of marked importance for GMP biology, including *Gfi1b, Meis1*, and *Runx2*. We suspect that these may be the most essential ER-HOXA9 targets, since they all have extensive gene regulatory potential. Several lines of evidence suggest that *Meis1* may play an especially critical role in driving the progenitor state transcriptional program. *Meis1* is highly expressed in progenitor cells and is sharply downregulated over the differentiation program (Figure 6A). Loss of MEIS1 may strongly affect ER-HOXA9 binding patterns, as the *Meis1* binding site is highly overrepresented in the promoters of genes that ER-HOXA9 fails to re-bind when reactivated after the commitment point (Figure 6B). Importantly, the *Meis1* locus itself appears to be under significant epigenetic control. It is among the most dramatically epigenetically inactivated loci in post-commitment point ER-HOXA9 cells, losing all five of its proximal H3K27ac peaks and all four of its proximal ATAC-seq peaks (Figure 6D and data not shown). Accordingly, while *Meis1* is a robust ER-HOXA9 target in the progenitor state, epigenetic inactivation renders it inaccessible after the commitment point, and its ER-HOXA9 binding site is among the most markedly reduced of all peaks in ER-HOXA9 ChIP-seq of post-vs. pre-commitment cells (Figures 6C and 6D). Consequently, *Meis1* downregulation is irreversible after the commitment point, and it remains silenced when ER-HOXA9 is reactivated after the differentiation point-of-no-return (Figure 6E)

**Figure 6.**
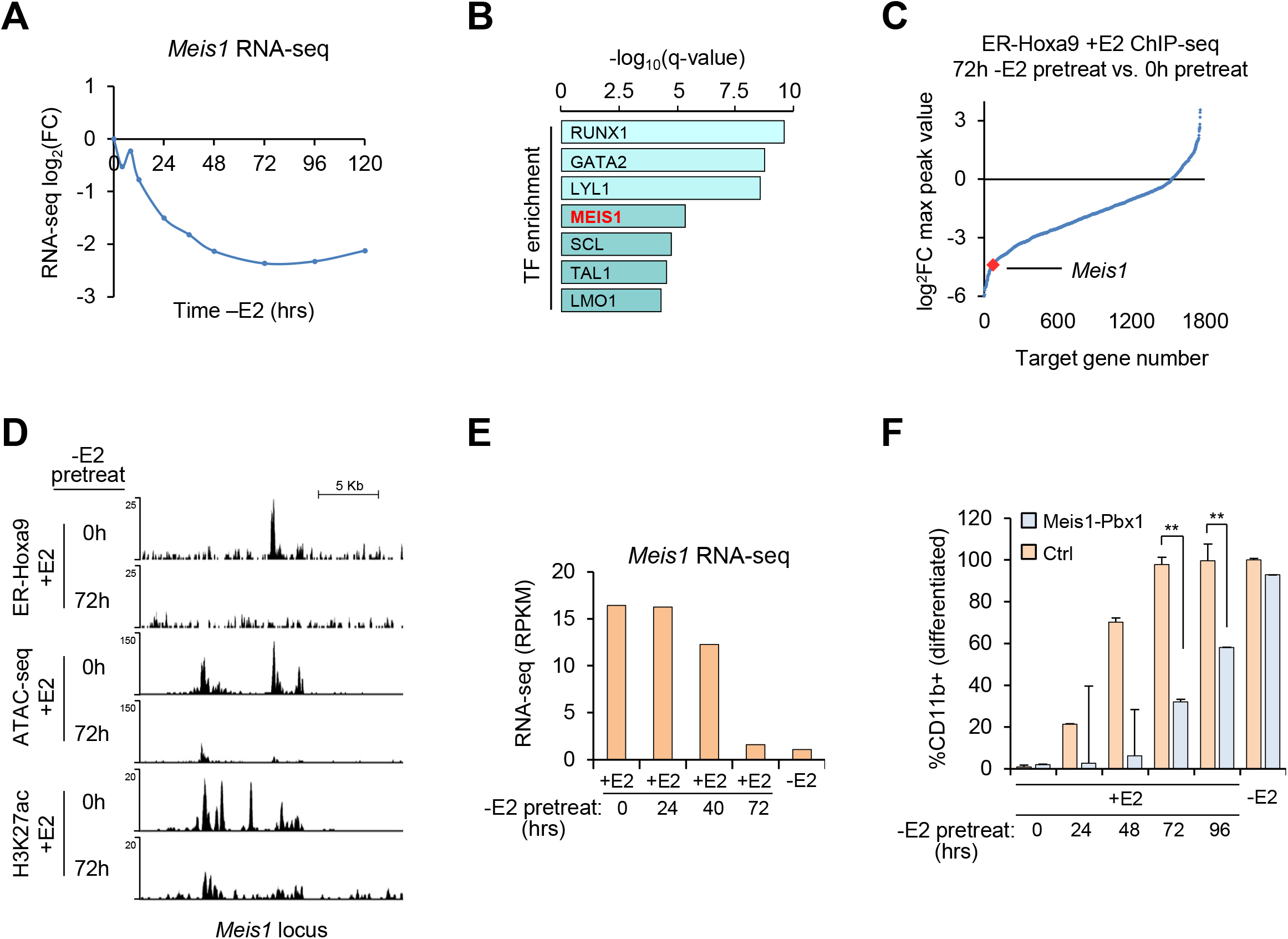
Re-expression of *Meis1* co-factor TF extends the pre-commitment window. A) Downregulation of *Meis1* over the -E2 differentiation time-course. B) Top ChIP-seq-based transcription factor target enrichments in the set of genes with the most reduced V5-ER-HOXA9-AM binding in cells cultured with vs. without 72-hour -E2 pretreatment. C) Plot of genes with proximal V5-ER-HOXA9-AM peaks, ordered by log2(fold-change) of maximum ChIP-seq peak value in cells cultured with vs. without 72-hour -E2 pretreatment. D) Tracks from V5-ER-HOXA9-AM ChIP-seq, H3K27ac ChIP-seq, and ATAC-seq of cells cultured with or without 72-hour -E2 pretreatment. E) Expression of *Meis1* in cells cultured +E2, 120 hours -E2, or +E2 after -E2 pretreatments. F) Live cell CD11b+ percentages of control or Meis1/Pbx1-overexpressing ER-Hoxb8 cells on Day 7 of commitment point assay. Cells were cultured +E2 for 7 days, -E2 for 7 days, or -E2 for the indicated durations followed by +E2 for the remainder of the experiment. **\***t-test p < 0.05; **t-test p < 0.01.

MEIS1 is well-known to heterodimerize with HOX9 and enhance its DNA-binding and trans-activation potential^37^. It follows that direct regulation of *Meis1* by HOXA9 creates a positive feedback loop that would be irreversibly broken by ER-HOXA9 inactivation and subsequent closing of *Meis1* regulatory sites. This suggests the possibility that, if *Meis1* could be expressed after the commitment point, this feedback loop would be restored, and ER-HOXA9 may have a greater ability to access and activate its progenitor state gene regulatory targets.

To test the role of MEIS1 in differentiation commitment, we derived new cell lines of the ER-Hoxb8 system harboring constitutive expression of *Meis1* and *Pbx1*. We used ER-Hoxb8 cells, as they more stably expressed exogenous *Meis1* than ER-HOXA9 cells. Constitutive expression of *Meis1* and *Pbx1* did not affect the ability of cells to differentiate upon inactivation of the ER-HOXB8 protein, as these cells upregulated CD11b along the same time course as control cells expressing ER-HOXB8 only (data not shown).

Commitment point assays were performed in which we inactivated ER-HOXB8 (E2 withdrawal) for varying periods of time before reactivation (E2 add-back). As expected, the majority of ER-HOXB8 cells could not be reverted to the progenitor state after 72h -E2, and effectively all cells were fully committed to terminal differentiation after 96h out of estrogen (Figure 6F). In contrast, though CD11b levels of live ER-HOXB8-MEIS1-PBX1 cells were near their maximum at 72-96 hours out of estrogen, the reactivation of ER-HOXB8 even at 96 hours -E2 induced significant portion of cells to decrease CD11b expression and to return toward the progenitor state (Figure 6F). In control ER-HOXB8 cells lacking *Meis1* expression, >99.9% of cells had terminally differentiated and died by this point, with only a fraction of a percent of outliers still alive and not terminally differentiated. These data suggest that co-expression of ER-HOXB8, MEIS1, and PBX1 endows ER-HOXB8 cells with the ability to go further into the myeloid differentiation program before irreversibly committing to differentiation.

### Chromatin accessibility and HOXA9 activity human and mouse myeloid differentiation programs *in vivo*

As the ER-HOXA9 model faithfully recapitulates the *in vivo* myeloid differentiation program^22^, we hypothesized that the key features of our model (Figure 7) would also be observed in epigenomic analyses of normal primary human and mouse myeloid cell populations. Accordingly, we analyzed ATAC-seq and RNA-seq data from sorted human and mouse GMPs and mature monocytes^38–40^ (neutrophil data were not available), as comparator cell types to the ER-HOXA9 cells in the progenitor and fully differentiated states. We first tested whether the *in vivo* progression from immature GMP to mature monocyte featured one of the hallmarks of our commitment point model – a dramatic global reduction in chromatin accessibility.

**Figure 7.**
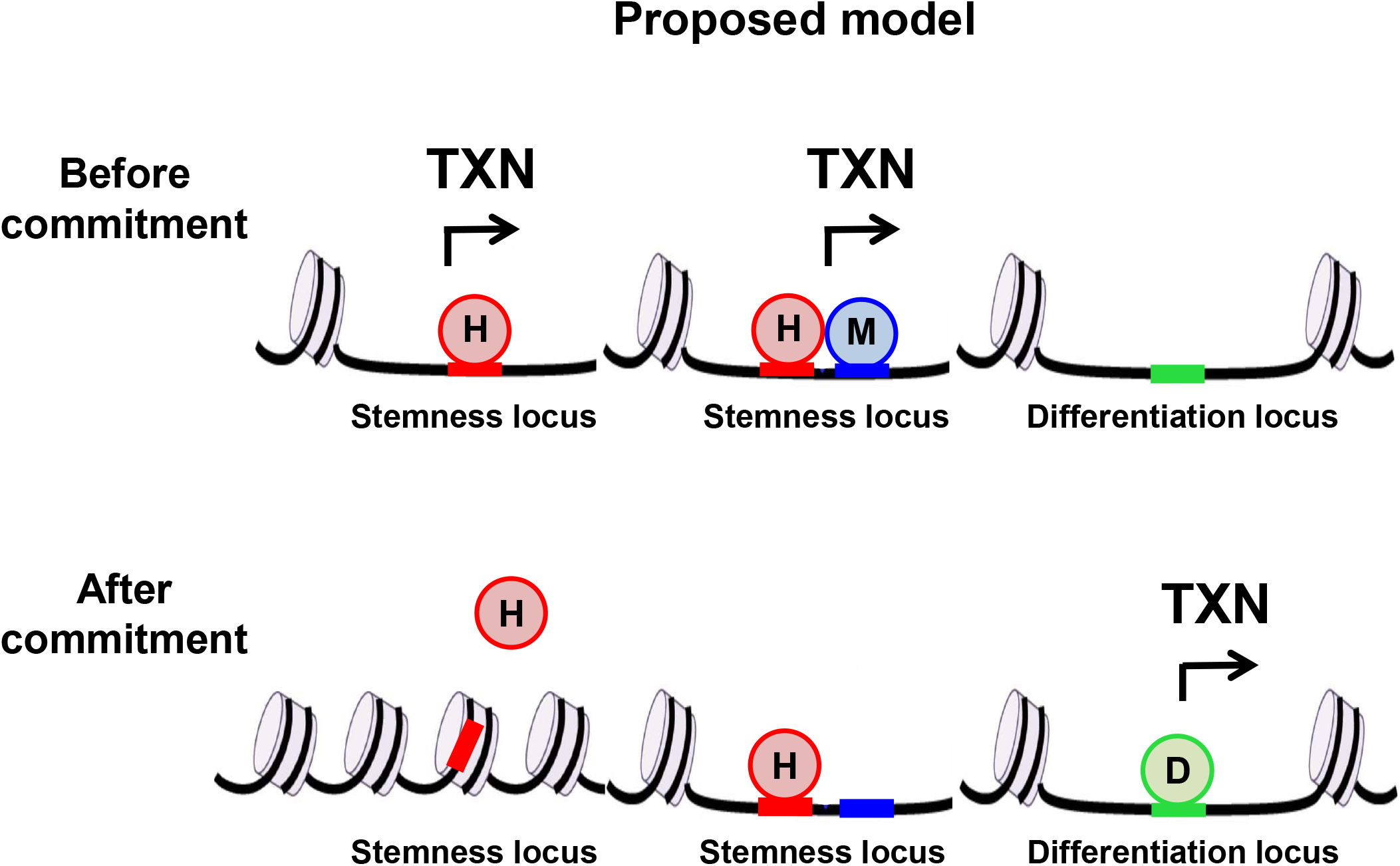
Model for molecular basis of point of no return. Chromatin focused model of TF expression, chromatin accessibility, and cell identity fate the pre-commitment (top) and post-commitment (bottom) states.

Consistent with our data, we found that 130,325 ATAC-seq peaks are lost or significantly reduced in the transition from human GMPs to monoctyes, while only 25,758 peaks (∼5-fold fewer) are gained or significantly increased (Figure S7A). Interrogating this program in a parallel dataset of sorted murine hematopoietic cells, we found that 141,465 peaks are lost/reduced in monocytes compared to GMPs, while only 35,432 peaks (∼4-fold fewer) are gained/increased (Figure S7A). We also investigated whether, as in ER-HOXA9 cells, regions of chromatin that are open in differentiated cells are already open in the progenitor state. In line with our model, the majority of regions of accessible chromatin in mouse monocytes are already accessible in GMPs (73,455 of 124,901 (59%); Figure S7B).

We then considered peak distribution trends in the myeloid differentiation program. As ER-HOXA9 cells differentiate, the percentage of promoter ATAC-seq peaks significantly increases, while the percentage of intron and intergenic peaks decreases (Figure 1E). Similarly, in human GMPs, 93% of ATAC-seq peaks that are lost/reduced in monocytes map to introns and intergenic regions, while only 2.6% map to promoters. Among peaks that are gained/increased in monocytes, only 60% map to introns and intergenic regions, while a striking 32% map to promoters (Figure S7C and Supplementary Table 7).

We next investigated putative HOXA9 targets and their transcriptional trends in the human dataset and compared them to our experimental data. We defined putative direct activation targets of ER-HOXA9 as genes that (1) have a proximal ER-HOXA9 ChIP-seq peak in the progenitor state, and (2) are significantly downregulated within the first 48 hours of the differentiation program – yielding 216 genes (Figure 4E). As the human datasets lack HOXA9 ChIP-seq, we defined the set of candidate direct activation targets of HOXA9 as genes that (1) had a HOXA9 consensus binding site within a GMP ATAC-seq peak that was lost/reduced in monocytes, and (2) were significantly transcriptionally downregulated in monocytes compared to GMPs - yielding 1,258 genes. Despite the differences in species (mouse vs. human), differentiated cell types (neutrophils vs. monocytes), datasets (with or without HOXA9 ChIP-seq) and contexts (*in vitro* vs. *in vivo*), we still observed a statistically significant overlap in the putative direct activation targets of HOXA9 in these two settings. Of the 216 direct activation targets in ER-HOXA9 cells, 31 – roughly 3-fold more than expected by chance - were conserved in human GMPs (p = 7.04 × 10-7, hypergeometric distribution). Among these conserved targets were several genes expected to be critical drivers of the ER-HOXA9 progenitor state, including *Flt3* and *Runx2* (Figure S7D).

Taken together, these data suggest that salient epigenomic features of our ER-HOXA9-based model of differentiation commitment are conserved *in vivo* in human hematopoietic cells.

## Discussion

### Models of cell fate commitment

Lineage choice and cellular differentiation programs proceed unidirectionally during development and in the maintenance of adult tissue. This process has been conceptualized in ‘Waddington’s Landscape’ as a progressive loss of pluripotency accompanying each cell identity decision^41^. The molecular underpinnings of cell identity barriers remain understudied. Repressive, self-perpetuating chromatin modifications such as H3K27me3^10^, H3K9me3^12^, and DNA methylation^11^ are proposed to establish these barriers by facilitating irreversible silencing of stem or progenitor cell expression programs during differentiation. However, these models are typically derived from correlative studies in embryonic stem cells^7–9^, and studies have not yet experimentally tested which of these chromatin features play a causal role in preventing de-differentiation.

### Identification of the “point of no return” in the myeloid differentiation program

Here we addressed the question of cell fate irreversibility using a tractable system of myeloid differentiation. In this model, conditional activity of the ER-HOXA9 homeobox transcription factor governs the differentiation of granulocyte-monocyte precursor (GMP) cells. GMPs with active ER-HOXA9 remain in an undifferentiated and self-renewing state; upon inactivation of ER-HOXA9 the cells initiate their myeloid differentiation program to terminally differentiated neutrophils. By inactivating and then reactivating ER-HOXA9 (via the removal and re-addition of estrogen), we mapped a window of commitment, prior to which the cells can revert to the self-renewing progenitor state, and beyond which they are destined to terminal differentiation.

While this program proceeds rapidly, the early stages (up to ∼40 hours) are reversible. However, after ∼72 hours, reactivation of ER-HOXA9 fails to revert cells to their progenitor state. Surprisingly, dynamics in promoter DNA methylation or H3K27me3 did not contribute to this cell identity commitment. Rather, we found that chromatin remodeling, loss of active regulatory sites, and co-factor silencing collectively form an epigenetic barrier to prevent ER-HOXA9 from re-establishing the progenitor-state transcriptome after the commitment point. This contrasts with proposed models of ESC differentiation commitment, in which repressive histone modifications and DNA methylation can irreversibly silence pluripotency factors such as *Oct-3/4* and prevent dedifferentiation^11,12^.

Our study focused on the choice of myeloid progenitor commitment from a self-renewing cell to a terminally differentiated effector cell. In this context, the “soft” epigenetic processes of chromatin accessibility and enhancer dynamics were sufficient to define a “point of no return.” We defined a role of the enhancer landscape in differentiation commitment, as the ER-HOXA9 protein is largely an enhancer-binding TF^34,35^. After inactivating ER-HOXA9, many of its target enhancers are also inactivated. Lacking active enhancers, the reactivated ER-HOXA9 protein – which is not thought to have pioneering factor activity at most of its binding targets^42,43^ - is unable to reactivate the majority of its original target genes. While enhancers are known to define cell identity, our work describes a new role of enhancer accessibility in preventing de-differentiation. This model does not preclude a role for DNA methylation, H3K27me3, or other chromatin features in differentiation commitment in other contexts, especially in early embryonic development.

The question remains as to how and why chromatin and enhancer accessibility is so dramatically reduced as the ER-HOXA9 cells differentiate. Epigenomic features can be “passively” lost via mitosis^44,45^. However, ER-HOXA9 cells exit the cell cycle within 72 hours after initiating differentiation^22^, and most losses in chromatin accessibility occur beyond this point (Figure 1D). A likely possibility is that the open chromatin at enhancers and promoters closes by default once transcription at the locus ceases. This would be consistent with models holding that the default state of chromatin is nucleosomal, and that regulatory loci are closed unless actively kept open^46,47^.

### Addressing Heterogeneity

As we followed the population of ER-HOXA9 cells during myeloid differentiation, we observed subtle cellular heterogeneity in the time required to reach the differentiation commitment point. One likely variable is the cell cycle status, as terminal differentiation programs are intimately linked to the cell cycle^48^, and it is possible that the “point of no return” occurs after a fixed number of cell cycles. In this case, the relatively broad commitment window (40 – 72 hours -E2) may be tightened in future experiments with synchronization of the cell cycle.

It is also possible that there is cell state heterogeneity in the starting population. Pluripotent cells are proposed to cycle through multiple pluripotency sub-states that may prime them for more rapid differentiation down particular lineages^49,50^. Whether GMPs cycle through analogous sub-states is an open question; single-cell transcriptomic and epigenomic approaches may be applied at several points along the differentiation program to address this question.

### Relevance to acute myeloid leukemia

Acute myeloid leukemia (AML) is characterized by hallmark differentiation arrest that maintains the leukemic blasts in a proliferative, self-renewing, progenitor-like state. “Differentiation therapy” with all-*trans* retinoic acid (ATRA) and arsenic trioxide offers curative therapy to patients with the promyelocytic AML subtype^51,52^. Inhibition of mutant IDH2 in another small subset of AML with IDH mutations may also prove effective by inducing differentiation^53–56^. Unfortunately, differentiation therapy has not been available for patients of other AML subtypes.

In addition to its use in studies of normal myeloid differentiation, the ER-HOXA9 model system also effectively models the differentiation block of AML. *Hoxa9* is upregulated in the majority (70%) of AMLs^57^ and is considered a critical driver of oncogenic differentiation arrest^37^. By delineating the mechanisms of differentiation irreversibility in the ER-HOXA9 model, our work aims to define the molecular processes underlying leukemic differentiation arrest and to identify those processes that may be amenable to therapeutic targeting in the development of new AML differentiation therapies.

Our model predicts that factors regulating chromatin remodeling and accessibility represent key vulnerabilities that AML cells need to overcome in order to avoid irreversible differentiation. Identification of such factors will be important to developing AML therapeutic strategies, and the ER-HOXA9 model of conditional differentiation arrest offers a tractable discovery system. Targeting differentiation-suppressive proteins or associated complex members may identify new approaches to differentiation therapy. Future work will determine the generalizability of this endeavor as well as the genetic and mutational backgrounds in which it might be most effective.

## Acknowledgments

We would like to acknowledge Katrina Maxcy for her help in sample preparation. We thank Aimee Iberg, Eric L. Greer, and Chris Lengner for helpful discussions. We thank Active Motif for their help with V5-tag and AM-tag chromatin immunoprecipitation. Flow cytometry instrumentation, sorting, and analysis were provided by the HSCI-CRM Flow Cytometry Core. We thank the Tufts University Core Facility for performing next-generation sequencing. We thank Jason Buenrostro and Vinay Kartha for helping with access to primary human and mouse hematopoietic cell ATAC-seq and RNA-seq datasets. M.A.B was supported by a Special Fellow Award from the Leukemia & Lymphoma Society (3353-15) and an NCI K22 award. D.B.S. was supported by a Scholar Award from the American Society of Hematology and an NCI K08 award. L.G. was supported by the Deutsche Forschungsgemeinschaft (DFG, German Research Foundation), EXC 2026, Cardio-Pulmonary Institute (CPI), Project ID 390649896. S.C. and K.H. were supported by the NIH grant 5R01HD058013-10, and SC is currently supported by the City of Hope/UCR biomedical research initiative (CUBRI) and UC cancer research coordinating committee (CRCC) seed grants. A.M. was supported by the NIH grant NIH 5P01GM099117 and the Max Planck Society. This work was also supported by grants to Y.S. from the Samuel Waxman Cancer Research Foundation, funds from Boston Children’s Hospital, and a grant to Y.S. and D.T.S. from the Harvard Epigenetics Initiative. This project was also partially supported by funds from the Ludwig Institute for Cancer Research to YS. YS is an American Cancer Society Research Professor.

## Author Contributions

M.A.B. and D.B.S. conceived and performed experiments, performed ChIP-seq and RNA-seq data analysis, interpreted all data sets, and wrote the manuscript.

L.G. performed the formal analysis of all ATAC-seq data, performed clustering of RNA-seq data, and assisted with ChIP-seq and RNAs-seq bioinformatics analyses.

M.Wu. worked with L.G. in all bioinformatics analyses. M.Wawer performed components of the RNA-seq analyses.

S.C. Performed the ATAC-seq experiments, intellectually contributed to overall study design, and assisted with data interpretation and manuscript writing.

R.P. and W.J. assisted with ChIP-seq experiments and sequencing library preparations.

J.R. and H.L. performed commitment point assays in additional ER-Hox cell systems

R.K. performed the RRBS methylation experiments and data analysis.

A.J. assisted with study design and data interpretation.

A.M. assisted with RRBS data analysis and interpretation.

K.H. assisted with ATAC-seq data analysis and interpretation.

Y.S. and D.T.S. helped with study design, data interpretation, and manuscript writing.

## Declaration of Interests

DBS is a co-founder and holds equity in Clear Creek Bio and SAFI Biosolutions. He is a consultant for Keros Therapeutics.

DTS is a co-founder and equity holder in Fate Therapeutics, Clear Creek Bio and LifeVault Bio; he is a director co-founder and equity holder in Magenta Therapeutics, he is a director and equity holder in Agios Pharmaceuticals and Editas Medicine, he is a consultant for FOG Pharma and VCanBio, a DSMB member for Alexion and a sponsored research recipient from Novartis.

YS is a co-founder and holds equity in Constellation Pharmaceuticals, Inc, Athelas Therapeutics, Inc and K36 Therapeutics, Inc. YS also holds equity in Imago Biosciences Inc, and is a consultant for Active Motif, Inc.

## STAR Methods

### CONTACT FOR REAGENT AND RESOURCE SHARING

Request for more information about this manuscript and any reagents or codes used therein will be fulfilled upon request to the Lead Contact, M. Andrés Blanco.

#### Cell culture

The ER-HOXA9 and ER-HOXB8 cell lines have been previously described in detail^22,23^. In brief, murine bone marrow was harvested by crushing the femur and tibia bones. The cells were filtered through a 40-micron filter and layered over a Ficoll-Paque-Plus gradient to collect live mononuclear cells. Cells were cultured for 48 hours in RPMI supplemented with 10% fetal bovine serum, penicillin/streptomycin, SCF (10 ng/ml), IL-3 (10 ng/ml) and IL-6 (10 ng/ml). Tissue culture-treated 12-well plates were pre-coated with human plasma fibronectin (1 ml per well, overnight at 37°C, at a final concentration of 10 µg/ml in phosphate buffered saline (PBS)). Fibronectin was aspirated prior to the addition of cells. At ∼48 hours, 2.5 × 10^5^ cells in a volume of 500 µl were transferred to each well. Polybrene was added and the cells were transduced via spinoculation (1000 g, 90 min, 22-degrees) with 1 ml of ecotropic retrovirus (MSCVneo-EE-ER-HOXA9; the EE denoting a GLU-GLU epitope tag or MSCVneo-HA-ER-HOXB8; the HA denoting a hemagluttinin epitope tag). The transduction volume was 1.5 ml with a polybrene concentration of 8 µg/ml. Following the transduction, 3 ml of fresh media was added to each well to dilute the polybrene to a less toxic concentration.

These GMP progenitors were maintained in RPMI supplemented with 10% fetal bovine serum, penicillin/streptomycin, and stem cell factor (SCF). The source of stem cell factor was conditioned media generated from a Chinese hamster ovary (CHO) cell line that stably secretes SCF. Conditioned medium was added at a final concentration of 1-2% (depending on the batch, final concentration of SCF approximately 100 ng/ml as measured by ELISA). Beta-estradiol (abbreviated E2, Sigma, E2758) was added to a final concentration of 0.5 µM from a 10 mM stock dissolved in 100% ethanol. The media was stable for at least 4 weeks when maintained at 4°C.

To inactivate ER-HOXA9 or ER-HOXB8, the cells were washed (2X PBS) free of beta-estradiol and re-plated in the same base media without estradiol.

#### Establishing the V5-Hoxa9-AM, V5-ER-HOXA9-AM, and ER-Hoxb8-Meis1-Pbx1 cell lines

Bone marrow mononuclear cells were isolated from female CD45.1STEM mice [https://pubmed.ncbi.nlm.nih.gov/27185283/] by Ficoll-Paque-Plus density gradient centrifugation and transduced as described above. The V5-HOXA9-AM, and ER-HOXB8-MEIS1-PBX1 constructs were cloned into the backbone of MSCVdeltaNEO (MSCVneo in which the PGK promoter and neomycin resistance cassette was removed by BglII + BamH1 digestion and ligation. This was done to decrease the size of the final MSCV construct and to allow for the generation of higher titer virus.

Gene synthesis and cloning of the constructs into the MSCVdeltaNEO backbone was done by VectorBuilder and ecotropic retrovirus generated by transient transfection of 293T cells using the Lipofectamine 2000 reagent.

#### Flow cytometry

Antibodies were purchased from BioLegend. Cells were suspended in FACS (fluorescence activated cell sorting) buffer (PBS + 2% FBS + 1 mM EDTA) and stained for 45 minutes at 4°C in the dark. 7-AAD or propidium iodide was included as a viability dye to help identify dead cells. Flow cytometry data was collected on either a BD FACSCalibur or BD LSR2 flow cytometer and analyzed using FlowJo software.

#### ChIP-seq

Cells were cross-linked with 1% formaldehyde at 37°C for 10 minutes, quenched with 125 mM glycine at room temperature for 5 minutes, and homogenized via douncing in swelling buffer. Nuclei were isolated via centrifugation and fragmented in sonication buffer by probe sonication. Insoluble material was removed via centrifugation and 500 ug chromatin (histone modification ChIPs) or 30 µg (V5-ER-HOXA9-AM) ChIPs was immunoprecipitated overnight at 4°C with 5-10 µg of antibody (for histone modification ChIPs) or 4 µg (for V5-ER-HOXA9-AM) ChIPs and 50 µl protein G magnetic dynabeads. Immunoprecipitated material was eluted by heating twice at 65°C for 5 minutes in 1% SDS buffer. Cross-links were reversed by incubating at 65°C overnight in 160 mM NaCl and 20 µg/ml RNase A. Samples were then treated with 200 µg/ml Proteinase K for two hours at 45°C and DNA recovered by phenol-chloroform extraction. 10 ng of DNA per sample was used for next generation sequencing library preparation using an NEBNext ChIP-seq Library Prep Master Mix Set for Illumina kit (New England Biolabs) as per manufacturer’s guidelines using AMPure XL beads for purification (Beckmann Coulter). 50 bp single end sequencing of DNA was performed on an Illumina HiSeq 2500 sequencer (histone modification ChIP-seq) or an Illumina NextSeq 500 sequencer (V5-ER-HOXA9-AM ChIP-seq). The following antibodies were used for chromatin immunoprecipitation: H3K4me3: Millipore CMA304; H3K27me3: Millipore 07-449; H3K27ac: Active Motif AM39135; V5: Abcam ab15828; AM: Active Motif 91111.

#### ChIP-seq data analysis

Reads were aligned to the genome using Bowtie 2.0^58^. Peaks were called using MACS or MACS 2.0^59^, depending on the sample, with mfold minimum set to 10 and FDR < 0.01. GO category enrichment analyses were performed with Genomic Regions Enrichment of Annotations Tool (GREAT)^60^ and EnrichR^61,62^, and motif enrichment analyses were performed with Analysis of Motif Enrichment (AME) from the MEME Suite 4.12.0^63^. Peaks were visualized using the UCSC Genome browser^64^. Superenhancer peaks were called as described previously^65^. Briefly, H3K27ac peaks were first called by using MACS2 and peaks within 12.5 kb of one another were stitched together as a super enhancer. ChIP-seq heat maps were generated using deepTools^66^ as follows. bamCompare was used to generate paired bigwig files using 50 bp bins, computeMatrix was used (parameters: -- beforeRegionStartLength 2000; --binSize 10; --sortUsing mean; --averageTypeBins mean) to generate scores for heat maps, and heat maps were generated with with plotHeatmap (parameters: –sortRegions descending order; – sortUsing mean; –averageTypeSummaryPlot mean). Genomic distributions of ChIP-seq peaks were generated using ChIPseeker^67^ with default parameters. For analysis of V5-ER-HOXA9-AM ChIP-seq data, 75-nt single-end sequence reads were mapped to the mm10 genome using the BWA algorithm with default settings. Reads that passed Illumina’s purity filter, had no more than 2 alignment mismatches, and mapped uniquely to the genome were de-duplicated and maintained for subsequent analysis using MACS for peak calling. For peak-to-gene mapping, peaks were assigned to the closest gene within a gene margin of 10,000 bp upstream to 10,000 bp downstream. If the peak was not within the margin for any genes, it was not given a gene association. Differential peak values for a given gene when comparing any two timepoints were defined as the log2 fold change between the fragment densities of the gene-associated peak interval at the two timepoints. For genes with multiple peaks within the gene margin, the peak interval with the highest fragment density was used.

#### ATAC-seq

To generate ATAC-seq libraries, 50,000 cells were used and libraries were constructed as previously described^68,69^. Briefly, cells were washed in PBS twice, counted and nuclei were isolated from 100,000 cells using 100 µl hypotonic buffer (10 mM Tris pH 7.4, 10 mM NaCl, 3 mM MgCl2, 0.1% NP40) to generate two independent transposition reactions. Nuclei were split in half and treated with 2.5 µl Tn5 Transposase (Illumina) for 30 mins. at 37°C. DNA from transposed nuclei was then isolated and PCR-amplified using barcoded Nextera primers (Illumina). Library QC was carried out using high sensitivity DNA biolanalyzer assay and qubit measurement and sequenced using paired-end sequencing (PE50) on the Illumina Hi-Seq 2500 platform.

#### ATAC-seq data analysis

Raw data were mapped to mouse reference genome version mm9 by BWA^70^. Potential PCR duplicates and ambiguously mapped reads were further filtered out for downstream analysis. Peaks were called by MACS 2.0^59^ with parameters callpeak --nomodel --extsize 200 --shift -100 -q 0.01. Consistency of peak calling results between replicates was examined using IDR analysis. IDR consistency threshold of 0.05 was applied to truncate the peak lists, peaks that passed the threshold were used for subsequent analysis. PCAs of ATAC-seq peak files were generated using ggplot2^71^ and ggfortify^72,73^, with MultiCovBed used to generate input files from peak bed files (default settings used for all parameters). For analysis of *in vivo* samples, ATAC-seq data were obtained from GSE74912 (human GMPs), direct transfer from Jason D. Buenrostro (human monocytes) and GSE143270 (mouse GMPs and monocytes). Briefly, raw fastq data were downloaded from SRA, reads passed QC were then mapped to hg19 and mm10 by BWA, and peaks were called by MACS2.

#### ATAC-seq motif enrichment analysis

ATAC-seq peak regions were analyzed for the occurrences of known motifs in HOMER motif databases^74^ using HOMER. -log10 (motif P-value) was used for the motif enrichment heatmap plot.

#### ATAC-seq time course clustering analysis

To obtain a reference peak set, ATAC-seq peaks of all time points were pooled and overlapping peaks regions (≥1bp) were merged into a single region which was the widest region that covers all merged regions. Raw reads were counted in the reference peaks set for each time point using Rsubread and differential analysis was performed using edgeR. Regions that showed no significant changes (|log2FC| > 2, FDR < 0.01) between any two time points were eliminated and remaining regions used for following clustering analysis. Fuzzy clustering cmeans^75^ was applied to recognize temporal patterns of the time course data.

#### RNA-seq

Normalized 10 timepoint, 120-hour RNA-seq data were published previously^22^. E2 withdrawal pretreatment and add-back RNA-seq datasets were analyzed similarly. Briefly, Reads were aligned to the mouse GRCm38 primary assembly extended with the sequence of eGFP (http://www.ebi.ac.uk/ena/data/view/AHK23750) using STAR^76^ and the GENCODE annotation M5 (www.gencodegenes.org). Differential expression analysis was performed with DESeq2 version 1.8.1^77^ in R version 3.2.1^78^. Significantly differentially expressed genes, when comparing any one timepoint to any other timpoint, were defined as genes with log2 fold change >1 and Benjamini Hochberg adjusted p-value < 0.05. Principal component analysis was performed in R and plotted using the ggplot2 package version 2.2.0^71^. Clustering analyses of RNA-seq datasets was performed as described for ATAC-seq clustering analyses. Heat maps were generated either using Heatmapper^79^ using average clustering and Euclidean distance metric, or using GSEA (below). For RNA-seq analysis of human *in vivo* samples, data were obtained from GSE74246, and DESeq2 was used to identify differential expression genes.

#### Gene Set Enrichment Analysis

Normalized time-course RNA-seq data were analyzed via GSEA 2.0^27,28^ after condensing the dataset to one value per gene by retaining the gene reading with the highest median RPKM value across all timepoints. Enrichment analyses were performed using 1,000 gene set permutations, a weighted enrichment statistic, the Signal2Noise ranking metric, and gene set minimum and maximum sizes of 15 and 500, respectively. Gene sets were obtained from the Molecular Signatures Database (MSigDB)^27,31,32^ or from ChIP-seq datasets generated in this study.

#### Reduced Representation Bisulfite Sequencing (RRBS)

Reduced Representation Bisulfite Sequencing (RRBS) Illumina libraries were prepared according to a standard gel-free pipeline^80^.

Reads were aligned to build mm9 of the mouse genome with MAQ^70^ in bisulfite alignment mode. For each CpG, the methylation level was computed as the number of reads with unconverted cytosines divided by the number of total reads covering that CpG. Mean methylation for each genomic feature was calculates as the mean methylation level of the CpGs within the feature weighted by the sequence coverage at each CpG, and only CpGs covered at 5x or higher were used in this calculation. Differentially methylation testing was performed by using methylation levels of CpGs covered at 5X or higher in a two-sample weighted coverage t-test. FDR q-values were calculated from the t-test p-values using the R q-value package^81^. Differentially methylated features were required to have a methylation difference of 0.2 at a FDR q-value threshold of 0.05.

#### RNA extraction

RNA was purified using an RNeasy kit (Qiagen) as per manufacturer’s instructions.

## Supplemental Figure Legends

**Figure S1. Gene Set Enrichment Analysis and GO analysis during myeloid differentiation. Related to Figure 1**. A) RNA-sequencing at 10 time points in biological duplicate following inactivation of ER-HOXA9 (upon removal of estradiol) demonstrates a graded, continuous progression of the transcriptome from progenitors to terminally differentiated neutrophils and macrophages over 120 hours. B) GSEA plots of most enriched (top plots) and most repressed (bottom panels) gene sets in RNA-seq ranked by expression of genes at 120 hours or 96 hours -E2 (differentiated cells) compared to +E2 (progenitor cells). C) Plots of GSEA normalized enrichment scores (NES) of gene sets over the -E2 differentiation time-course. Left - selected gene sets shown in (B). Right – sets of genes with promoters containing the indicated TF binding sites. D) Quantification of total number of TF or miRNA motifs enriched at *p* < 0.01 in promoters of up-regulated or down-regulated genes over the time-course. All RNA-seq data represent the average of RNA-seq experiments performed in biological duplicate at each time point.

**Figure S2. Global overviews of ATAC-seq profiling of the 120-hour ER-HOXA9 differentiation time-course. Related to Figure 1**. A) Principal component analysis (PCA) of ATAC-seq of ER-HOXA9 cells across the differentiation program. B) ATAC-seq peak overlap between progenitor state (+E2) and terminally differentiated (-E2) ER-HOXA9 cells. C) Tracks showing ATAC-seq signals at the *Flt3* (left) *Alcam* (middle) and *S100a9* (right) loci across the differentiation program, representing the three main chromatin accessibility trends observed. Expression of each gene across the differentiation program is shown beneath the gene’s ATAC-seq tracks. D) TF binding site motif enrichments in ATAC-seq peaks of ER-HOXA9 cells in the progenitor state (+E2) and the terminally differentiated state (120h -E2).

**Figure S3. Identifying the point-of-no-return in alternative clones and stability of DNA methylation during myeloid differentiation. Related to Figure 2**. To demonstrate that the window of differentiation commitment was not isolated to an individual clone of cells, differentiation commitment assays were performed as shown in Figure 2B, only in a different ER-HOXA9 clone (A), and in the complementary ER-Hoxb8 differentiation arrest model (B). Live cell GFP+ cell percentages are shown in each case. In (B), GFP was quantified on day 5 (left) and again from the same samples at day 7 (right), to confirm that the cells do not revert to the progenitor state if given more time. Beyond day 7 (which is 4 days after +E2 add-back to 72-hour -E2 pretreatment samples), differentiated cell populations were almost entirely dead, precluding extension of the assay further. All results shown represent the average + standard deviation of experiments performed at least in biological triplicate. C) Clustering of samples by Pearson correlation of global DNA methylation patterns, as assayed by reduced representation bisulfite sequencing (RRBS). Samples were cultured +E2, -E2 for 120 hours, or +E2 after varying durations of -E2 pretreatments. Assays were performed in biological duplicate. D) RNA-seq log2(fold change) of genes with increasing or decreasing promoter DNA methylation in 120-hour -E2 (differentiated) cells vs. +E2 (progenitor state) cells. E) DNA methylation dynamics in regions containing H3K27ac ChIP-seq peaks following the same treatments as in (C).

**Figure S4. H3K27 methylation does not influence myeloid differentiation gene expression programs. Related to Figure 2**. A) Heat map of normalized H3K27me3 ChIP-seq read counts at all peak regions centered +/- 2Kb from the peak center in progenitor state (+E2) or terminally differentiated (-E2) ER-HOXA9 cells. B) Distribution of H3K27me3 ChIP-seq peaks according to genome annotation from progenitor (+E2) and terminally differentiated (-E2) ER-HOXA9 cells. C) Tracks showing H3K27me3 ChIP-seq signal at the *Hoxa* locus in progenitor cells (0h -E2), terminally differentiated cells (120h -E2), and cells cultured +E2 after 40h or 72h -E2 pretreatment. D) Violin plots of expression distribution of all genes losing (left plot) or gaining (right plot) H3K27me3 peaks in terminally differentiated (120h -E2) compared to progenitor state (+E2) ER-HOXA9 cells. Expression distributions of the same genes are shown in progenitor and terminally differentiated cells in each plot. p-values represent t-test results. E) Aggregate numbers of H3K27me3 ChIP-seq peaks proximal to genes in the “Brown Myeloid Cell Development Up” and “Brown Myeloid Cell Development Down” gene sets in progenitor (+E2) and terminally differentiated (120h -E2) ER-HOXA9 cells.

**Figure S5. Global analyses of ATAC-seq analyses from commitment point assays. Related to Figure 3**. A) Clustering of ATAC-seq data from ER-HOXA9 cells cultured +E2, -E2, or +E2 after -E2 pretreatments identifies 7 main patterns of chromatin accessibility dynamics. B) ATAC-seq tracks showing chromatin accessibility of 6 gene loci in ER-HOXA9 cells cultured in the progenitor state (0 h –E2 pretreatment), the differentiated state (-E2) or after 40 or 72 hours of –E2 pretreatment prior to E2 add-back.

**Figure S6. ChIP-seq of wild type (non-ER-tagged) V5-Hoxa9-AM with V5 and AM antibodies. Related to Figure 4**. A) Correlation of read tags from V5 and AM antibody ChIP-seq of V5-Hoxa9-AM. r indicates Pearson correlation coefficient. B) Heat map of merged peak regions (left, -/+ 5 Kb from peak center) or promoters (right, +/-5 Kb from TSS) from ChIP-seq of V5-Hoxa9-AM with V5 and AM antibodies. C) Distribution of peaks according to genome annotation from ChIP-seq of V5-Hoxa9-AM with AM and V5 antibodies. D) MEME enrichment of known DNA motifs within peaks from ChIP-seq of V5-ER-HOXA9-AM with AM and V5 antibodies. E) (Top) Overlap of top 100 most enriched GO biological processes between AM and V5 antibody ChIP-seq of V5-Hoxa9-AM. (Bottom) Overlap of genes with proximal peaks in ChIP-seq of V5-HOXA9-AM with AM and V5 antibodies. F) Plots of average normalized read density from ChIP-seq of V5-Hoxa9-AM with AM and V5 antibodies. Region plotted is -/+ 2 Kb from the TSS

**Figure S7. *in vivo* Chromatin accessibility and transcriptional trends in GMPs and monocytes from humans and mice**. **Related to Figure 5**. A) Number of ATAC-seq peaks lost/decayed or gained/enhanced in human and mouse monocytes compared to GMPs. B) ATAC-seq peak overlap between mouse GMPs (+E2) and monocytes (-E2). C) Number of ATAC-seq peaks that are lost/decayed or gained/increased in human monocytes compared to GMPs stratified by associated genome feature. D) Tracks showing ATAC-seq and RNA-seq trends in human GMPs and monocytes at the *Flt3* (top) and *Runx2* (bottom) loci. Hoxa9 binding motifs within peaks that are present in GMPs and absent in monocytes are highlighted.

## Supplemental Table Titles

Table S1. GSEA MSigDB C2 and C5 results from 120 hour -E2 RNA-seq time course.

Table S2. p-values from heat map in Figure 1F showing TF binding motif enrichment in accessible chromatin regions over the ER-HOXA9 differentiation program.

Table S3. Biological Process and Interpro Domain enrichments of H3K27me3 ChIP-seq results.

Table S4. Biological Process and Molecular Function enrichments of HOXA9 ChIP-seq results.

Table S5. V5-ER-HOXA9-AM ChIP-seq target genes lists.

Table S6. p-values from heat map in Figure 5G showing TF binding motif enrichment in regions of overlap between progenitor state ER-HOXA9 ChIP-seq and ATAC-seq peaks. Motif enrichments are shown for cells in the progenitor state (+E2), terminally differentiated state (-) E2, and cultured +E2 after 72 hour –E2 pretreatment.

Table S7. Number of ATAC-seq peaks lost/decayed or gained/increased in human monocytes compared to GMPs, stratified by associated genome feature.

## References

1. Murata, K. et al. Ascl2-Dependent Cell Dedifferentiation Drives Regeneration of Ablated Intestinal Stem Cells. Cell Stem Cell 26, 377-390.e6 (2020).

2. Schwitalla, S. et al. Intestinal tumorigenesis initiated by dedifferentiation and acquisition of stem-cell-like properties. Cell 152, 25–38 (2013).

3. Tata, P. R. et al. Dedifferentiation of committed epithelial cells into stem cells in vivo. Nature 503, 218–223 (2013).

4. Tetteh, P. W. et al. Replacement of Lost Lgr5-Positive Stem Cells through Plasticity of Their Enterocyte-Lineage Daughters. Cell Stem Cell 18, 203–213 (2016).

5. Doulatov, S., Notta, F., Laurenti, E. & Dick, J. E. Hematopoiesis: a human perspective. Cell Stem Cell 10, 120–136 (2012).

6. Nichols, J. & Smith, A. Pluripotency in the embryo and in culture. Cold Spring Harb Perspect Biol 4, a008128 (2012).

7. Gifford, C. A. et al. Transcriptional and epigenetic dynamics during specification of human embryonic stem cells. Cell 153, 1149–1163 (2013).

8. Suelves, M., Carrió, E., Núñez-Álvarez, Y. & Peinado, M. A. DNA methylation dynamics in cellular commitment and differentiation. Brief Funct Genomics 15, 443–453 (2016).

9. Xie, W. et al. Epigenomic analysis of multilineage differentiation of human embryonic stem cells. Cell 153, 1134–1148 (2013).

10. Lu, T. T.-H. et al. The Polycomb-Dependent Epigenome Controls β Cell Dysfunction, Dedifferentiation, and Diabetes. Cell Metab 27, 1294-1308.e7 (2018).

11. Feldman, N. et al. G9a-mediated irreversible epigenetic inactivation of Oct-3/4 during early embryogenesis. Nature Cell Biology 8, 188–194 (2006).

12. Nicetto, D. & Zaret, K. Role of H3K9me3 Heterochromatin in Cell Identity Establishment and Maintenance. Curr Opin Genet Dev 55, 1–10 (2019).

13. Bostick, M. et al. UHRF1 plays a role in maintaining DNA methylation in mammalian cells. Science 317, 1760–1764 (2007).

14. Hansen, K. H. et al. A model for transmission of the H3K27me3 epigenetic mark. Nat Cell Biol 10, 1291– 1300 (2008).

15. Sharif, J. et al. The SRA protein Np95 mediates epigenetic inheritance by recruiting Dnmt1 to methylated DNA. Nature 450, 908–912 (2007).

16. Zhu, B. & Reinberg, D. Epigenetic inheritance: Uncontested? Cell Research 21, 435–441 (2011).

17. Takahashi, K. & Yamanaka, S. Induction of pluripotent stem cells from mouse embryonic and adult fibroblast cultures by defined factors. Cell 126, 663–676 (2006).

18. Soufi, A. et al. Pioneer transcription factors target partial DNA motifs on nucleosomes to initiate reprogramming. Cell 161, 555–568 (2015).

19. Soufi, A., Donahue, G. & Zaret, K. S. Facilitators and impediments of the pluripotency reprogramming factors’ initial engagement with the genome. Cell 151, 994–1004 (2012).

20. Stricker, S. H., Köferle, A. & Beck, S. From profiles to function in epigenomics. Nat Rev Genet 18, 51–66 (2017).

21. Gunne-Braden, A. et al. GATA3 Mediates a Fast, Irreversible Commitment to BMP4-Driven Differentiation in Human Embryonic Stem Cells. Cell Stem Cell 26, 693-706.e9 (2020).

22. Sykes, D. B. et al. Inhibition of Dihydroorotate Dehydrogenase Overcomes Differentiation Blockade in Acute Myeloid Leukemia. Cell 167, 171-186.e15 (2016).

23. Wang, G. G. et al. Quantitative production of macrophages or neutrophils ex vivo using conditional Hoxb8. Nature Methods 3, 287–293 (2006).

24. Lawrence, H. J. et al. Mice bearing a targeted interruption of the homeobox gene HOXA9 have defects in myeloid, erythroid, and lymphoid hematopoiesis. Blood 89, 1922–1930 (1997).

25. Golub, T. R. et al. Molecular classification of cancer: class discovery and class prediction by gene expression monitoring. Science 286, 531–537 (1999).

26. Faust, N., Varas, F., Kelly, L. M., Heck, S. & Graf, T. Insertion of enhanced green fluorescent protein into the lysozyme gene creates mice with green fluorescent granulocytes and macrophages. Blood 96, 719–726 (2000).

27. Subramanian, A. et al. Gene set enrichment analysis: a knowledge-based approach for interpreting genome-wide expression profiles. Proc Natl Acad Sci U S A 102, 15545–15550 (2005).

28. Mootha, V. K. et al. PGC-1α-responsive genes involved in oxidative phosphorylation are coordinately downregulated in human diabetes. Nature Genetics 34, 267–273 (2003).

29. Meissner, A. et al. Reduced representation bisulfite sequencing for comparative high-resolution DNA methylation analysis. Nucleic Acids Res 33, 5868–5877 (2005).

30. Lam, K. et al. Hmga2 is a direct target gene of RUNX1 and regulates expansion of myeloid progenitors in mice. Blood 124, 2203–2212 (2014).

31. Liberzon, A. et al. Molecular signatures database (MSigDB) 3.0. Bioinformatics 27, 1739–1740 (2011).

32. Liberzon, A. et al. The Molecular Signatures Database (MSigDB) hallmark gene set collection. Cell Syst 1, 417–425 (2015).

33. Goldberg, L. et al. Genome-scale expression and transcription factor binding profiles reveal therapeutic targets in transgenic ERG myeloid leukemia. Blood 122, 2694–2703 (2013).

34. Huang, Y. et al. Identification and characterization of Hoxa9 binding sites in hematopoietic cells. Blood 119, 388–398 (2012).

35. Sun, Y. et al. HOXA9 Reprograms the Enhancer Landscape to Promote Leukemogenesis. Cancer Cell 34, 643-658.e5 (2018).

36. Yu, H. & Gerstein, M. Genomic analysis of the hierarchical structure of regulatory networks. PNAS 103, 14724–14731 (2006).

37. Collins, C. T. & Hess, J. L. Deregulation of the HOXA9/MEIS1 axis in acute leukemia. Curr Opin Hematol 23, 354–361 (2016).

38. Corces, M. R. et al. Lineage-specific and single-cell chromatin accessibility charts human hematopoiesis and leukemia evolution. Nat Genet 48, 1193–1203 (2016).

39. Lal, A. et al. Deep learning-based enhancement of epigenomics data with AtacWorks. Nat Commun 12, 1507 (2021).

40. Xiang, G. et al. An integrative view of the regulatory and transcriptional landscapes in mouse hematopoiesis. Genome Res 30, 472–484 (2020).

41. Waddington. The Strategy of the Genes. (London: George Allen & Unwin). (George Allen & Unwin, 1957).

42. Porcelli, D., Fischer, B., Russell, S. & White, R. Chromatin accessibility plays a key role in selective targeting of Hox proteins. Genome Biol 20, 115 (2019).

43. Choe, S.-K., Ladam, F. & Sagerström, C. G. TALE factors poise promoters for activation by Hox proteins. Dev Cell 28, 203–211 (2014).

44. Margueron, R. & Reinberg, D. Chromatin structure and the inheritance of epigenetic information. Nat Rev Genet 11, 285–296 (2010).

45. Margueron, R. et al. Role of the polycomb protein EED in the propagation of repressive histone marks. Nature 461, 762–767 (2009).

46. Escobar, T. M. et al. Active and Repressed Chromatin Domains Exhibit Distinct Nucleosome Segregation during DNA Replication. Cell 179, 953-963.e11 (2019).

47. Becker, P. B. & Workman, J. L. Nucleosome Remodeling and Epigenetics. Cold Spring Harb Perspect Biol 5, (2013).

48. Kueh, H. Y., Champhekar, A., Nutt, S. L., Elowitz, M. B. & Rothenberg, E. V. Positive Feedback Between PU.1 and the Cell Cycle Controls Myeloid Differentiation. Science 341, 670–673 (2013).

49. Chambers, I. et al. Nanog safeguards pluripotency and mediates germline development. Nature 450, 1230– 1234 (2007).

50. Rodriguez-Terrones, D. et al. A molecular roadmap for the emergence of early-embryonic-like cells in culture. Nat Genet 50, 106–119 (2018).

51. Huang, M. E. et al. All-trans retinoic acid with or without low dose cytosine arabinoside in acute promyelocytic leukemia. Report of 6 cases. Chin Med J (Engl) 100, 949–953 (1987).

52. Shen, Z. X. et al. Use of arsenic trioxide (As2O3) in the treatment of acute promyelocytic leukemia (APL): II. Clinical efficacy and pharmacokinetics in relapsed patients. Blood 89, 3354–3360 (1997).

53. Losman, J.-A. et al. (R)-2-hydroxyglutarate is sufficient to promote leukemogenesis and its effects are reversible. Science 339, 1621–1625 (2013).

54. Rohle, D. et al. An inhibitor of mutant IDH1 delays growth and promotes differentiation of glioma cells. Science 340, 626–630 (2013).

55. Wang, F. et al. Targeted inhibition of mutant IDH2 in leukemia cells induces cellular differentiation. Science (New York, N.Y.) 340, 622—626 (2013).

56. Yen, K. et al. AG-221, a First-in-Class Therapy Targeting Acute Myeloid Leukemia Harboring Oncogenic IDH2 Mutations. Cancer Discov 7, 478–493 (2017).

57. Golub, T. R. et al. Molecular classification of cancer: class discovery and class prediction by gene expression monitoring. Science 286, 531–537 (1999).

58. Langmead, B. & Salzberg, S. L. Fast gapped-read alignment with Bowtie 2. Nat Methods 9, 357–359 (2012).

59. Zhang, Y. et al. Model-based Analysis of ChIP-Seq (MACS). Genome Biology 9, R137 (2008).

60. McLean, C. Y. et al. GREAT improves functional interpretation of cis-regulatory regions. Nat Biotechnol 28, 495–501 (2010).

61. Chen, E. Y. et al. Enrichr: interactive and collaborative HTML5 gene list enrichment analysis tool. BMC Bioinformatics 14, 128 (2013).

62. Kuleshov, M. V. et al. Enrichr: a comprehensive gene set enrichment analysis web server 2016 update. Nucleic Acids Res 44, W90–97 (2016).

63. McLeay, R. C. & Bailey, T. L. Motif Enrichment Analysis: a unified framework and an evaluation on ChIP data. BMC Bioinformatics 11, 165 (2010).

64. Kent, W. J. et al. The human genome browser at UCSC. Genome Res 12, 996–1006 (2002).

65. Whyte, W. A. et al. Master transcription factors and mediator establish super-enhancers at key cell identity genes. Cell 153, 307–319 (2013).

66. Ramírez, F., Dündar, F., Diehl, S., Grüning, B. A. & Manke, T. deepTools: a flexible platform for exploring deep-sequencing data. Nucleic Acids Res 42, W187–191 (2014).

67. Yu, G., Wang, L.-G. & He, Q.-Y. ChIPseeker: an R/Bioconductor package for ChIP peak annotation, comparison and visualization. Bioinformatics 31, 2382–2383 (2015).

68. Buenrostro, J. D., Giresi, P. G., Zaba, L. C., Chang, H. Y. & Greenleaf, W. J. Transposition of native chromatin for fast and sensitive epigenomic profiling of open chromatin, DNA-binding proteins and nucleosome position. Nat Methods 10, 1213–1218 (2013).

69. Cheloufi, S. et al. The histone chaperone CAF-1 safeguards somatic cell identity. Nature 528, 218–224 (2015).

70. Li, H., Ruan, J. & Durbin, R. Mapping short DNA sequencing reads and calling variants using mapping quality scores. Genome Res 18, 1851–1858 (2008).

71. Wickham, H. ggplot2: Elegant Graphics for Data Analysis. (Springer-Verlag, 2009).

72. Horikoshi, M. & Tang, Y. ggfortify: Data Visualization Tools for Statistical Analysis Results. (2018).

73. Tang, Y., Horikoshi, M., & W Li. ggfortify: Unified Interface to Visualize Statistical Result of Popular R Packages. The R Journal 8, (2016).

74. Heinz, S. et al. Simple combinations of lineage-determining transcription factors prime cis-regulatory elements required for macrophage and B cell identities. Mol Cell 38, 576–589 (2010).

75. Futschik, M. E. & Carlisle, B. Noise-robust soft clustering of gene expression time-course data. J Bioinform Comput Biol 3, 965–988 (2005).

76. Dobin, A. et al. STAR: ultrafast universal RNA-seq aligner. Bioinformatics 29, 15–21 (2013).

77. Love, M. I., Huber, W. & Anders, S. Moderated estimation of fold change and dispersion for RNA-seq data with DESeq2. Genome Biol 15, 550 (2014).

78. Team, R. C. R: A Language and Environment for Statistical Computing. In R Foundation for Statistical Computing (Vienna, Austria). (2005).

79. Babicki, S. et al. Heatmapper: web-enabled heat mapping for all. Nucleic Acids Res 44, W147–153 (2016).

80. Boyle, P. et al. Gel-free multiplexed reduced representation bisulfite sequencing for large-scale DNA methylation profiling. Genome Biol 13, R92 (2012).

81. Storey, J. D. & Tibshirani, R. Statistical significance for genomewide studies. Proc Natl Acad Sci U S A 100, 9440–9445 (2003).

