## Supplementary figures and images for "Chromatin state barriers enforce an irreversible mammalian cell fate decision"

### Supplementary Information

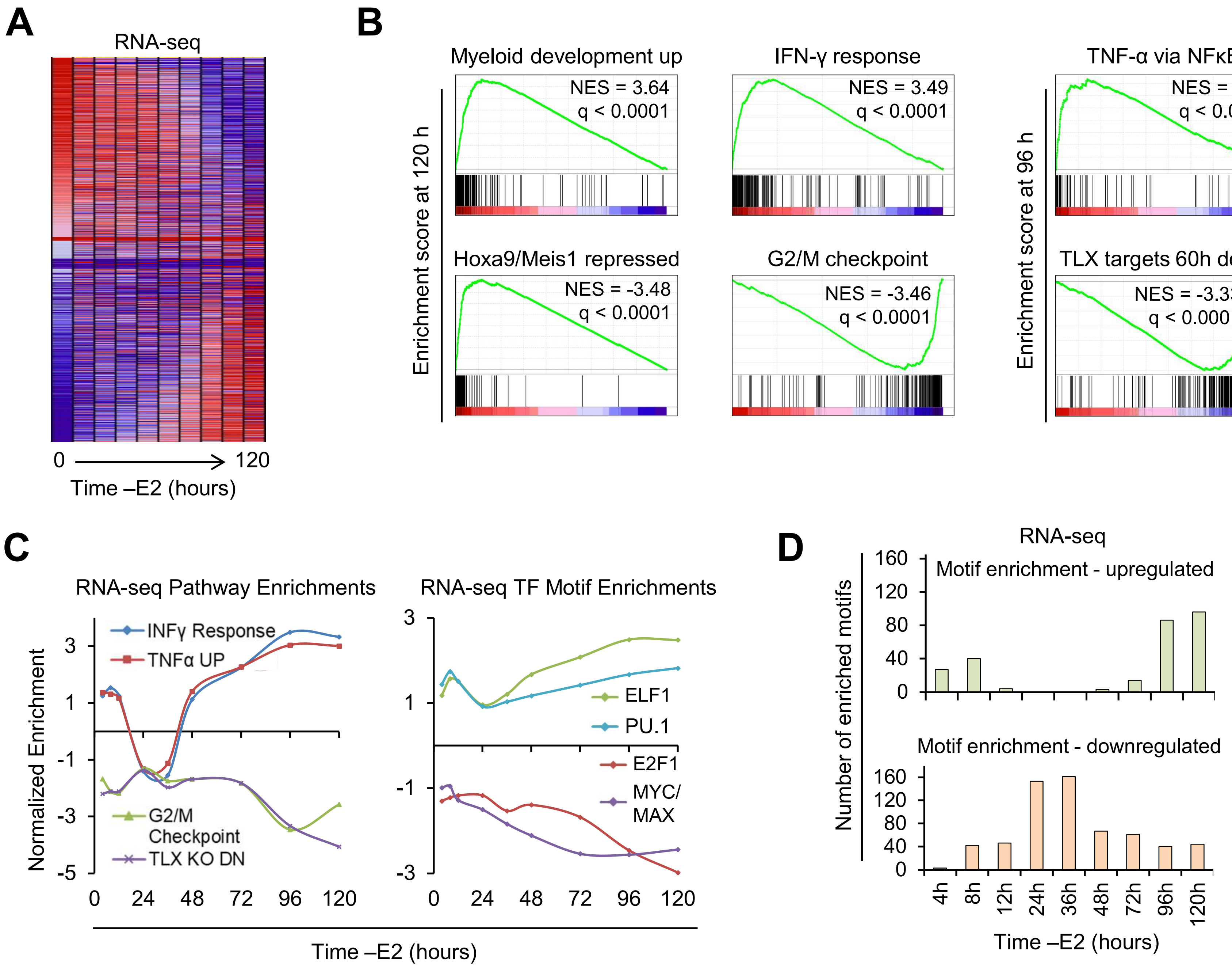

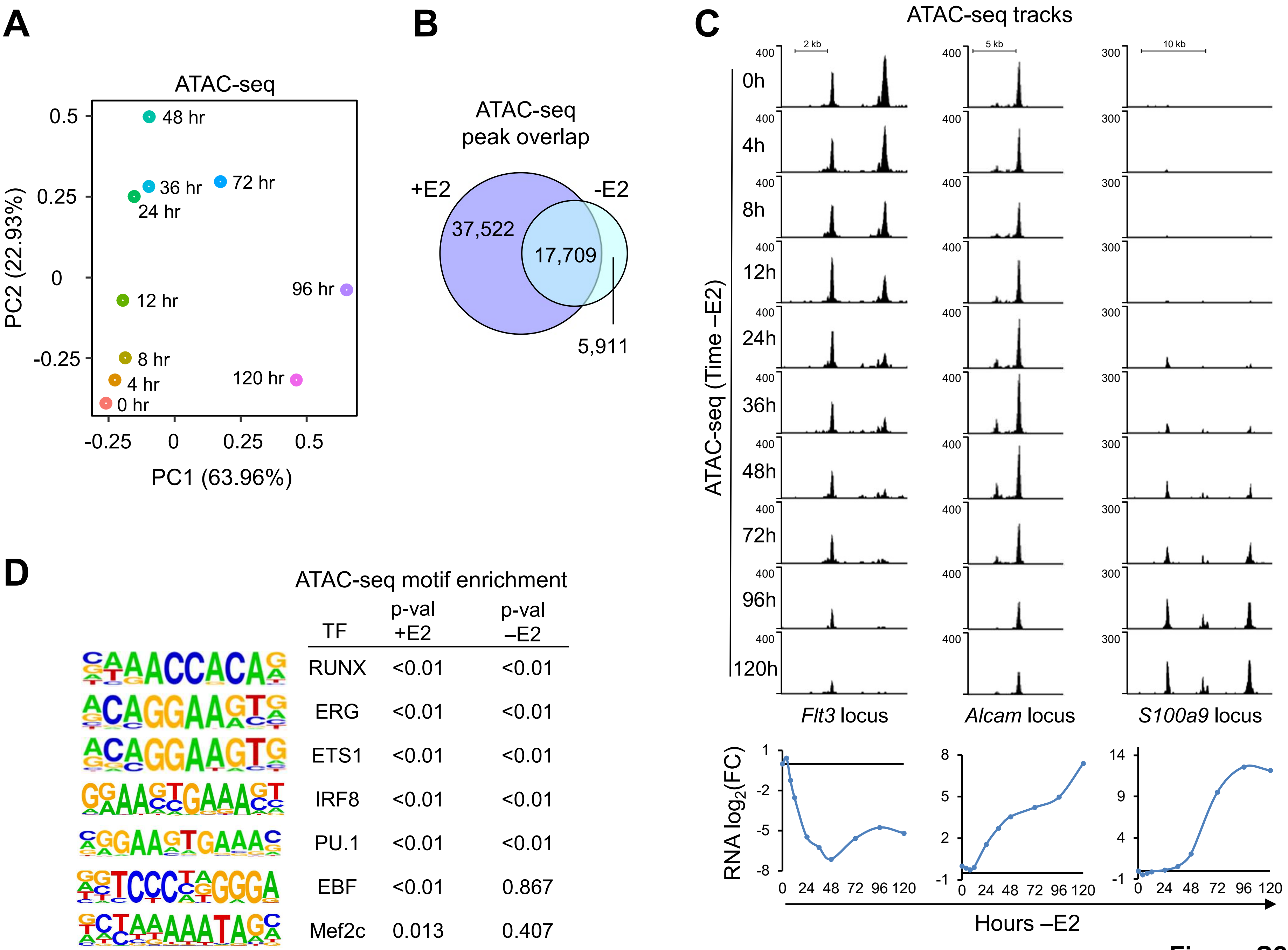

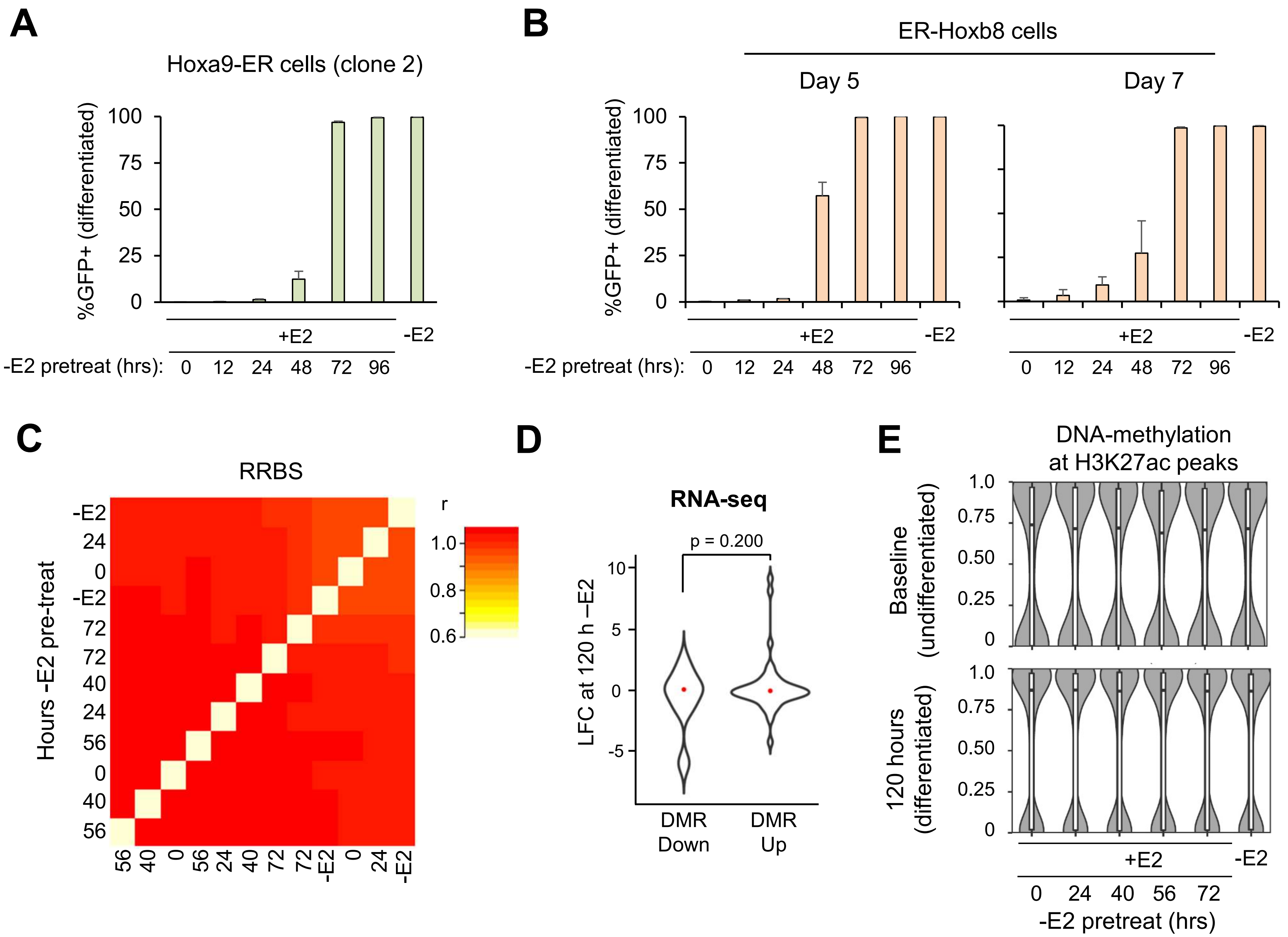

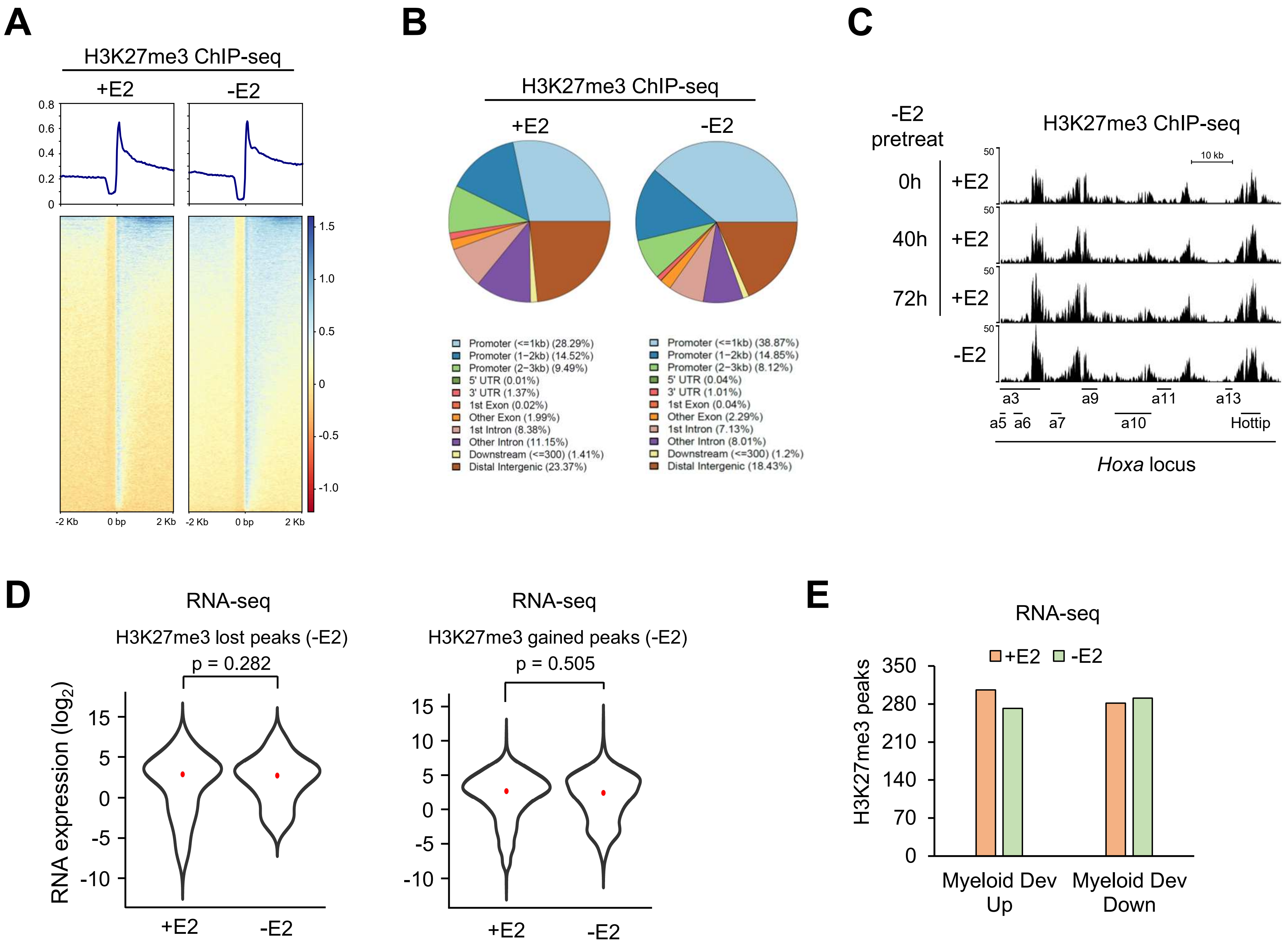

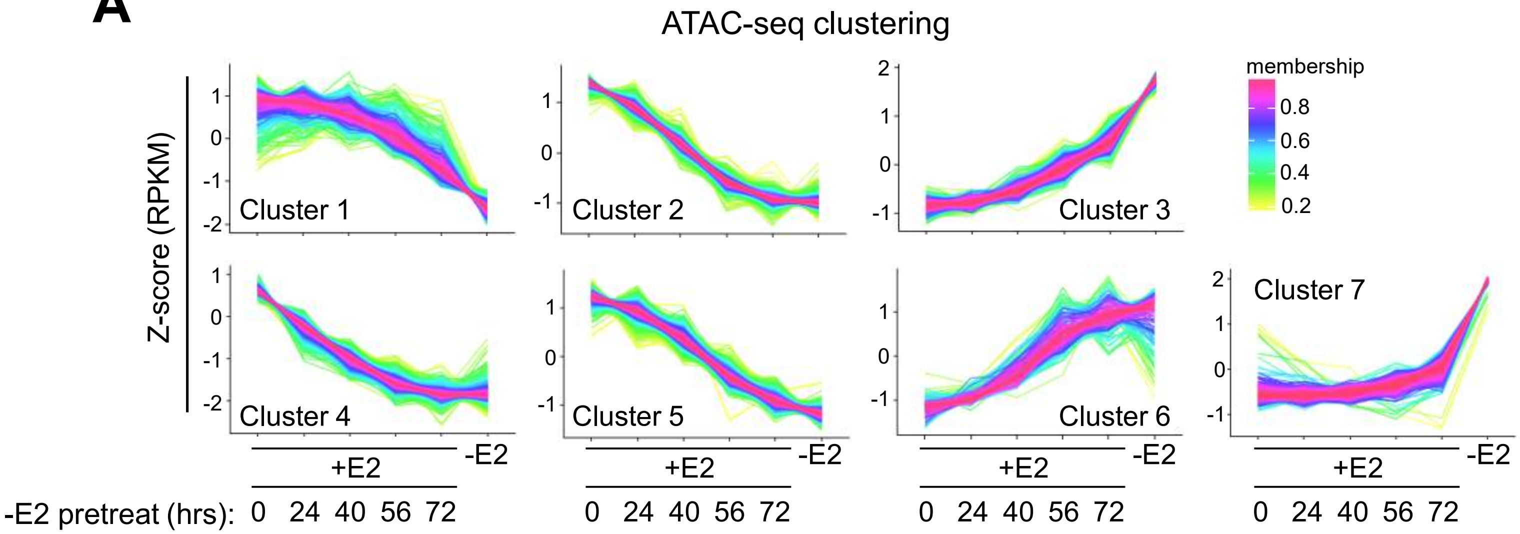

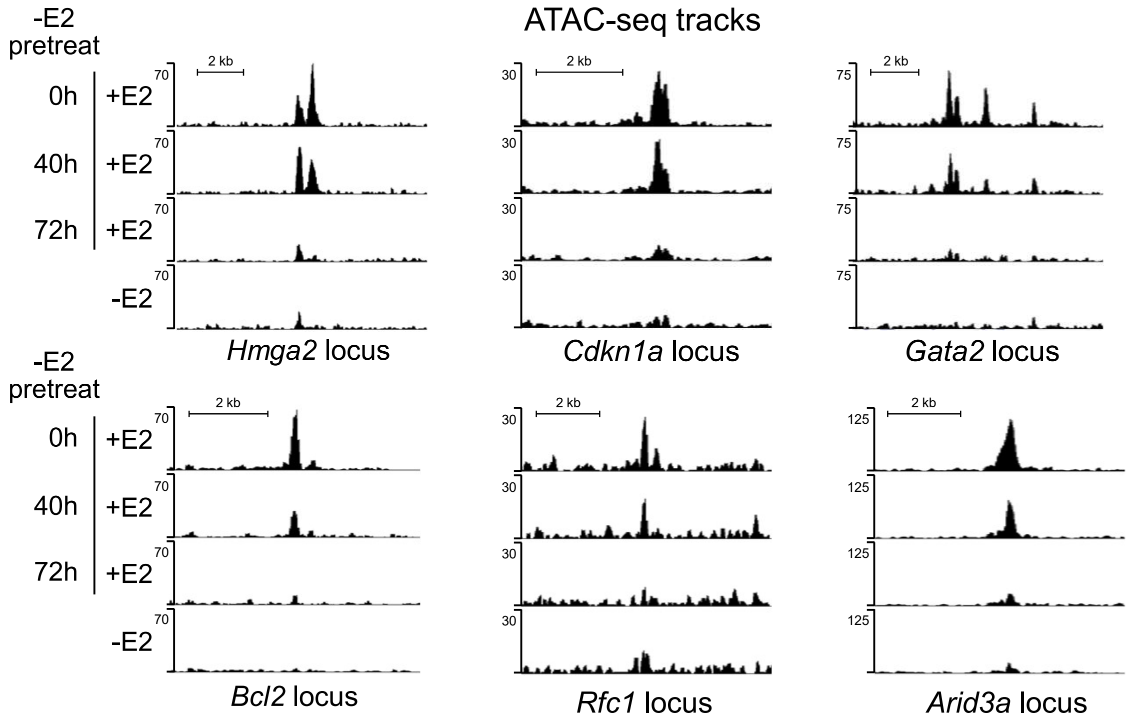

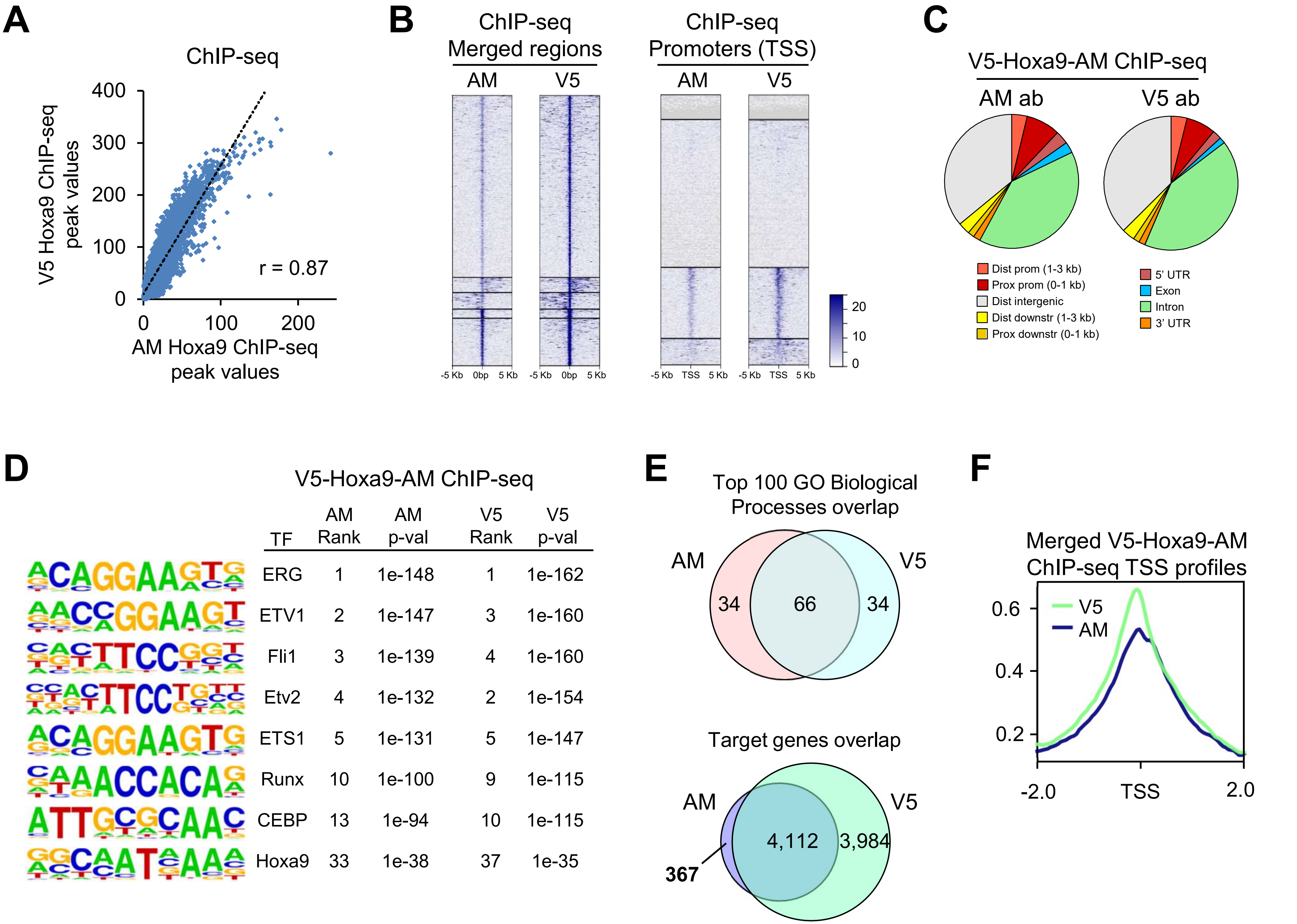


| TF | AM Rank | AM p-val | V5 Rank | V5 p-val |
|---|---|---|---|---|
| ERG | 1 | 1e-148 | 1 | 1e-162 |
| ETV1 | 2 | 1e-147 | 3 | 1e-160 |
| Fli1 | 3 | 1e-139 | 4 | 1e-160 |
| Etv2 | 4 | 1e-132 | 2 | 1e-154 |
| ETS1 | 5 | 1e-131 | 5 | 1e-147 |
| Runx | 10 | 1e-100 | 9 | 1e-115 |
| CEBP | 13 | 1e-94 | 10 | 1e-115 |
| Hoxa9 | 33 | 1e-38 | 37 | 1e-35 |

**Figure S6**

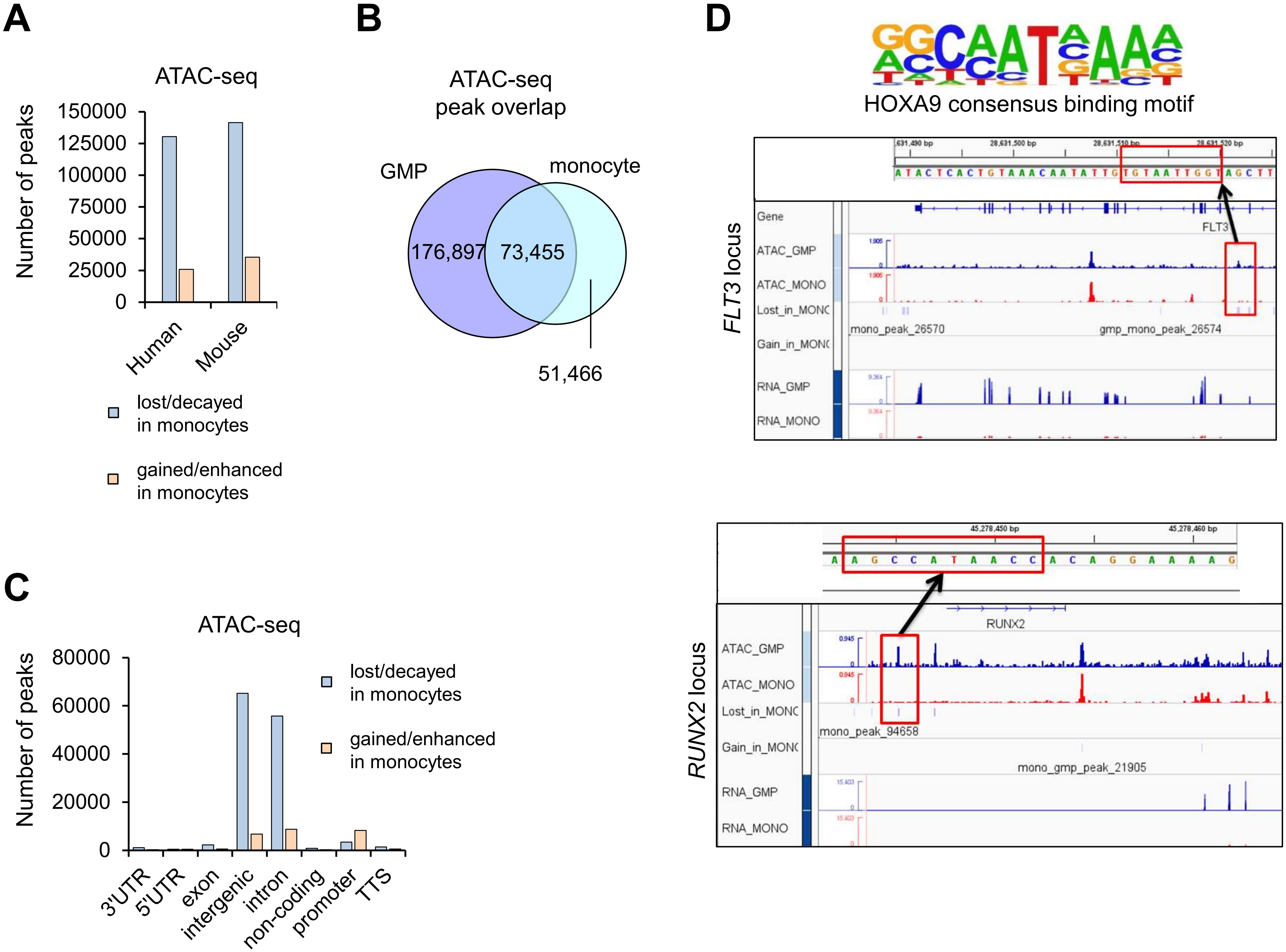
